# Antimicrobial resistance level and conjugation permissiveness shape plasmid distribution in clinical enterobacteria

**DOI:** 10.1101/2023.04.03.535338

**Authors:** Aida Alonso-del Valle, Laura Toribio-Celestino, Anna Quirant, Carles Tardio Pi, Javier DelaFuente, Rafael Canton, Eduardo Rocha, Carles Ubeda, Rafael Peña-Miller, Alvaro San Millan

**Affiliations:** Centro Nacional de Biotecnología (CNB), CSIC. Madrid, Spain; Fundación para el Fomento de la Investigación Sanitaria y Biomédica de la Comunitat Valenciana - FISABIO, Valencia, Spain; Centro de Ciencias Genómicas, Universidad Nacional Autónoma de México, Cuernavaca, Mexico; Instituto de Investigaciones en Matemáticas Aplicadas y en Sistemas, Unidad Académica Yucatán, Universidad Nacional Autónoma de México, Yucatán, México; Servicio de Microbiología. Hospital Universitario Ramón y Cajal-IRYCIS. Madrid, Spain; Centro de Investigación Biológica en Red de Enfermedades Infecciosas (CIBERINFEC), Instituto de Salud Carlos III, Madrid, Spain; Institut Pasteur, Université de Paris Cité, CNRS UMR3525, Microbial Evolutionary Genomics, Paris, France; Centro de Investigación Biológica en Red de Epidemiología y Salud Pública (CIBERESP), Instituto de Salud Carlos III, Madrid, Spain

**Author notes:** Contributed equally (Aida Alonso-del Valle, Laura Toribio-Celestino). Correspondence and request for materials should be addressed to Aida Alonso-del Valle & Alvaro San Millan.

## Abstract

Conjugative plasmids play a key role in the dissemination of antimicrobial resistance (AMR) genes across bacterial pathogens. AMR plasmids are widespread in clinical settings, but their distribution is not random, and certain associations between plasmids and bacterial clones are particularly successful. For example, the globally spread carbapenem resistance plasmid pOXA-48 can use a wide range of enterobacterial species as hosts, but it is usually associated with a small number of specific *Klebsiella pneumoniae* clones. These successful associations represent an important threat for hospitalized patients. However, knowledge remains limited about the factors determining AMR plasmid distribution in clinically relevant bacteria. Here, we combined *in vitro* and *in vivo* experimental approaches to analyze pOXA-48-associated AMR levels and conjugation dynamics in a collection of wild type enterobacterial strains isolated from hospitalized patients. Our results reveal significant variability in these traits across different bacterial hosts, with *Klebsiella* spp. strains showing higher pOXA-48-mediated AMR and conjugation frequencies than *Escherichia coli* strains. Using experimentally determined parameters, we developed a simple mathematical model to interrogate the contribution of AMR levels and conjugation permissiveness to plasmid distribution in bacterial communities. The simulations revealed that a small subset of clones, combining high AMR levels and conjugation permissiveness, play a critical role in stabilizing the plasmid in different polyclonal microbial communities. These results help to explain the preferential association of plasmid pOXA-48 with *K. pneumoniae* clones in clinical settings. More generally, our study reveals that species- and strain-specific variability in plasmid-associated phenotypes shape AMR evolution in clinically relevant bacterial communities.

**Significance statement:** Conjugative plasmids disseminate AMR genes across bacterial pathogens. Understanding the rules governing plasmid dynamics in bacterial communities is therefore crucial to controlling the global AMR crisis. In this study, we analyzed the dynamics of an AMR plasmid of great clinical relevance, pOXA-48, in a collection of wild type bacteria recovered from hospitalized patients. We reported a high degree of variability in two key plasmid-associated phenotypes, AMR level and conjugation ability, across the collection of clinical bacteria. Using simulations based on the experimental results, we studied how successful associations between AMR plasmids and clinical strains can arise in bacterial communities. Our results revealed that accounting for variability in plasmid-associated phenotypes help to understand the evolution of AMR in clinical settings.

## Introduction

Plasmids shape bacterial ecology and evolution by spreading accessory genes across populations. Plasmids can disseminate both vertically and horizontally. Vertical transmission is coupled to the division of the bacterial host, whereas horizontal transmission is mediated mainly by conjugation (the direct cell-to-cell transfer of plasmids through a bridge-like connection) (1). Vertical plasmid transmission is favored in the presence of environmental stresses that select for plasmid-encoded genes, but in the absence of plasmid-specific selection, vertical transmission is usually hindered by the fitness costs that plasmids produce in their bacterial hosts (2, 3). Therefore, plasmids’ interests can align or conflict with those of their bacterial hosts depending on the environmental conditions, leading to complex eco-evolutionary dynamics (4–6). Horizontal plasmid transmission can in principle make up for a deficit in vertical transmission, although conjugation rates vary considerably depending on the plasmid type (7, 8). The fate of plasmids in bacterial populations is thus determined by the interplay between horizontal and vertical transmission dynamics (and their evolution) (9–12). Multiple theoretical and experimental studies have investigated plasmid dynamics in clonal bacterial populations (examples can be found in 7, 8, 13–17). Other studies have analyzed the variability of plasmid-associated phenotypes across the diversity of wild type bacteria that plasmids encounter in natural communities (18–25). However, information remains very limited about how this variability may affect plasmid distribution in complex microbiota.

A dramatic example of the ability of plasmids to fuel bacterial evolution is the central role they play in the spread of antimicrobial resistance (AMR) in clinical pathogens, which is one of the most urgent public health threats facing humanity (26, 27). One particularly concerning group of drug resistant pathogens is carbapenemase-producing enterobacteria (order *Enterobacterales*), which appear on the WHO “priority pathogens” list (28, 29). Carbapenemases are β-lactamase enzymes able to degrade carbapenem antibiotics and are mainly acquired through conjugative plasmids (26). pOXA-48-like plasmids (from here on pOXA-48) constitute one of the most clinically important groups of carbapenemase-producing plasmids (30). pOXA-48 is a broad-host-range conjugative plasmid from the plasmid taxonomic unit L/M (31) that encodes the OXA-48 carbapenemase and is disseminated across enterobacteria worldwide (30). Crucially, like most AMR plasmids, pOXA-48 is not randomly distributed across bacterial hosts. Although pOXA-48 has a broad potential host range, it is strongly associated with *Klebsiella pneumoniae* and is especially prevalent in high-risk clones of specific sequence-types (ST), such as ST11, ST307 or ST101 (32, 33). In a previous study, we proposed that this bias in host distribution could be explained in part by pOXA-48-associated fitness costs in the absence of antibiotics (34). We analyzed the distribution of pOXA-48 fitness effects across a collection of *Escherichia coli* and *Klebsiella* spp. clinical strains. Our results revealed that pOXA-48 produced a wide distribution of fitness effects across these hosts, but these effects could not explain overall plasmid distribution, especially at the species level (34).

In the present study, we investigate how other key plasmid-associated phenotypes determine pOXA-48 distribution across clinical enterobacteria. We determine pOXA-48-associated AMR level and conjugation permissiveness in a collection of multidrug resistant *E. coli* and *Klebsiella* spp. strains isolated from the gut microbiota of hospitalized patients. Through a combination of *in vitro* and *in vivo* experimental approaches, we demonstrate that (i) pOXA-48 confers a higher level of AMR in *Klebsiella* spp., and that (ii) *Klebsiella* spp. strains are, on average, more permissive than *E. coli* strains to pOXA-48 acquisition through conjugation. Finally, we integrate these new experimentally determined parameters with our previous fitness data in a simple population dynamics model. We use this model to interrogate the distribution of plasmid pOXA-48 in complex bacterial communities under different antibiotic concentrations.

## Results

### Experimental model system

In this study, we used a collection of multidrug-resistant clinical enterobacterial strains isolated from the gut microbiota of patients at a large hospital in Madrid, Spain (Ramon y Cajal University Hospital). We focused on the two most prevalent species associated with pOXA-48 in this hospital, *K. pneumoniae* and *E. coli* (32). Since *K. quasipneumoniae* and *K. variicola* (two species previously misidentified as *K. pneumoniae*) are also associated with pOXA-48 in the hospital (32), we included strains from these species in the study. From now on we use *Klebsiella* spp. to refer to the strains in our collection belonging to these three *Klebsiella* species.

To analyze pOXA-48 transmission dynamics it is necessary to work with pOXA-48-free strains, which are able to receive the plasmid through conjugation. Therefore, we selected and sequenced the genomes of 25 *Escherichia coli* and 25 *Klebsiella* spp. multidrug resistant strains that did not carry this plasmid, and that were representative of the phylogenetic diversity of these species in our hospital (Fig. S10 and Table S2, collection partially overlapping with the one in ref. (34), see Methods). Although these strains were pOXA-48-naive, they were identified as ecologically compatible with pOXA-48 because they were obtained from patients on hospital wards where pOXA-48-carrying enterobacteria were commonly isolated during the same period (32, 34).

### AMR level conferred by pOXA-48 in wild type enterobacteria

To characterize the fitness advantage associated with pOXA-48 in the presence of antibiotics, we determined the impact of plasmid acquisition on the level of AMR. pOXA-48 was introduced by conjugation into the pOXA-48-naive enterobacteria collection. Four *E. coli* strains and one *K. pneumoniae* strain were unable to acquire the plasmid, and whole genome sequencing of the remaining transconjugants revealed pOXA-48 mutations in six *E. coli* and six *Klebsiella* spp. transconjugants. After excluding these strains, we used the remaining 15 *E. coli* and 18 *Klebsiella* spp. isogenic strains pairs with or without pOXA-48 plasmid for this analysis (Fig. S10). For each of the transconjugant (TC) and plasmid-free (PF) clones, we determined the minimal inhibitory concentration (MIC) of 4 clinically important β-lactam antibiotics: amoxicillin in combination with the β-lactamase inhibitor clavulanic acid (AMC) and the carbapenems ertapenem (ERT), imipenem (IMP), and meropenem (MER) (Fig. 1A). Compared with *E. coli*, *Klebsiella* spp. strains showed higher intrinsic resistance to all four antibiotics (Wilcoxon rank sum test: AMC, *W* = 47.5, *P* = 0.001; ERT, *W* = 14.5, *P* = 8.68 x 10^-6^; IMP, *W* = 15, *P* = 5.16 x 10^-6^; MER, *W* = 56, *P* = 0.002), as well as higher resistance level after plasmid acquisition (Wilcoxon rank sum test: AMC, *W* = 27, *P* = 8.94 x 10^-5^; ERT, *W* = 82, *P* = 0.031; IMP, *W* = 29.5, *P* = 4.79 x 10^-5^; MER, *W* = 54, *P* = 0.002).

**Fig. 1.**
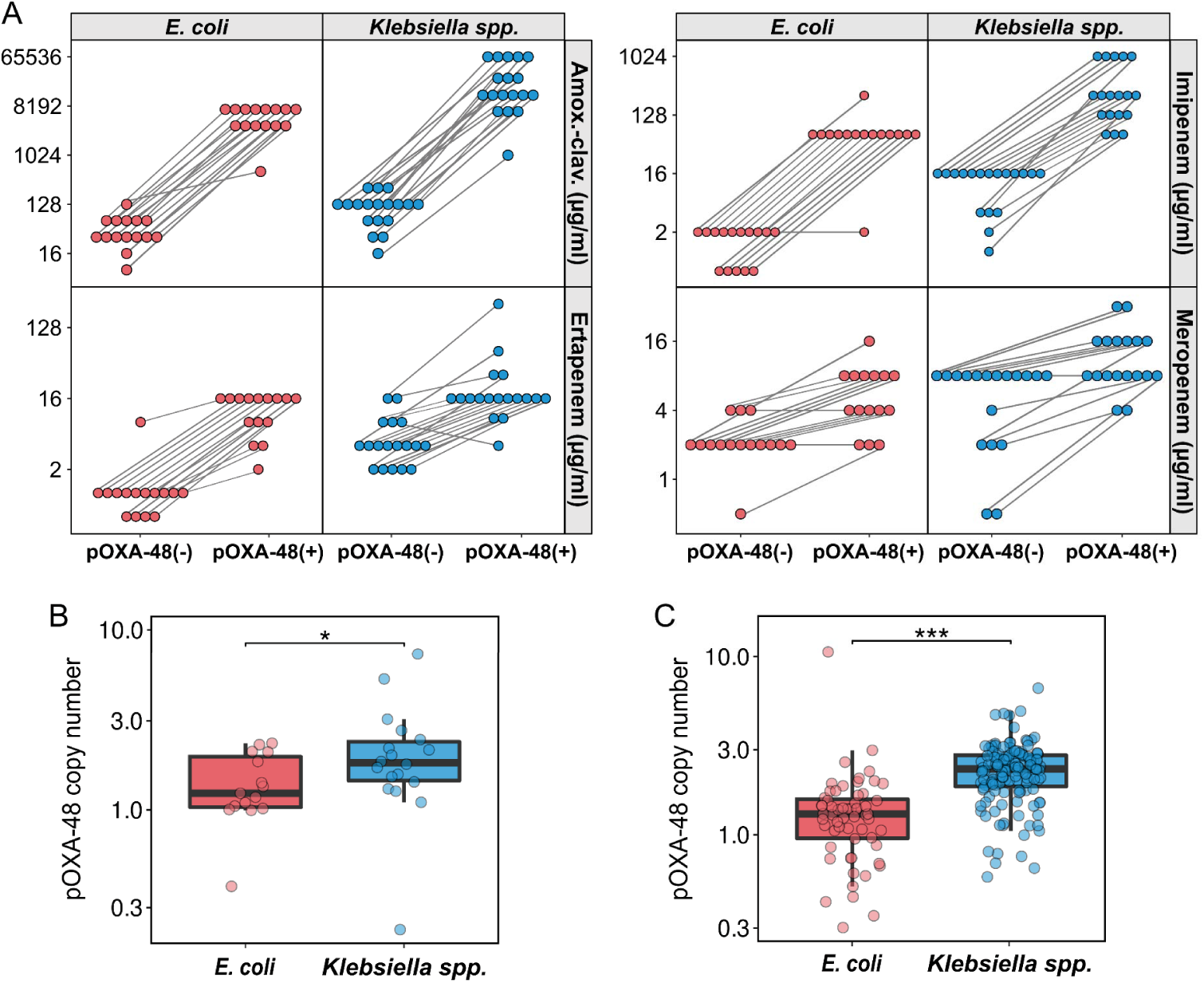
Resistance levels and copy number of pOXA-48 in wild type enterobacteria. pOXA-48 confers higher resistance levels and has a higher PCN in *Klebsiella* spp. than in *E. coli.* (A) MIC (mg/L) of amoxicillin/clavulanic acid, ertapenem, imipenem and meropenem for the plasmid-free/plasmid-carrying strain pairs (median of three biological replicates). (B) pOXA-48 plasmid copy number (PCN) of the *E. coli* (n = 15) and *Klebsiella* spp. (n = 18) transconjugant strains. PCN was estimated from sequencing data as the ratio of plasmid/chromosome median coverage (see Methods). Horizontal lines inside boxes mark median values, the upper and lower hinges correspond to the 25th and 75th percentiles, and whiskers extend to 1.5 times the interquartile range. Dots indicate individual PCN values. To aid visualization, the y axis is represented in log_10_ scale with no value transformation. (C) pOXA-48 PCN in *E. coli* (n = 59) and *Klebsiella* spp. (n = 140) pOXA-48-carrying wild-type clinical strains. Boxplots are structured as in B. \**P* < 0.05, \*\**P* < 0.01, \*\*\**P* < 0.001.

The diverse genomic background and AMR gene content of the strains under study made it difficult to elucidate a genetic origin for the higher AMR levels reached by pOXA-48-carrying *Klebsiella* spp. (Fig. S11). However, a prominent influence on plasmid-mediated AMR level is plasmid copy number (PCN). AMR level commonly escalates with PCN for AMR mechanisms showing strong gene dosage effects, such as carbapenemases (35, 36). We used the genome sequencing data to estimate pOXA-48 PCN in the strains under study (Fig. 1B). The results showed that PCN was higher in *Klebsiella* spp. (mean = 2.21, SD = 1.53) than in *E. coli* strains (mean = 1.41, SD = 0.55) (Wilcoxon rank-sum test, *W* = 76, *P* = 0.03). We then compared pOXA-48 PCN and AMR levels for the antibiotic to which pOXA-48 confers the highest level of resistance: AMC (average fold-change in MIC associated to pOXA-48 presence = 244.7, Fig. S12). The final AMC resistance level correlated positively with PCN (Spearman’s correlation: *R* = 0.5, *P* = 0.003; Fig. S13), supporting the idea that the higher PCN is at least partly responsible for the elevated AMR level in *Klebsiella*. spp. Finally, to investigate if the differences in PCN were also present in strains naturally carrying pOXA-48, we used the same genomic approach to analyze PCN in a recently characterized large collection of 200 pOXA-48-carrying clinical isolates from the same hospital. In line with the results from the transconjugant strains, PCN was higher in *Klebsiella* spp. (mean = 2.35, SD = 0.88) than in *E. coli* (mean = 1.46, SD = 1.33) (Fig. 1C; Wilcoxon rank-sum test, *W* = 1271, *P* = 1.32 x 10^-14^).

### Conjugation frequency varies across clinical strains

To analyze pOXA-48 horizontal transmission dynamics, we determined the ability of clinical strains to acquire the plasmid by conjugation (“conjugation permissiveness”). For recipients, we used a subset of 10 *Klebsiella spp.* and 10 *E. coli* strains representative of the genetic diversity of the collection (Fig. S10). For donors, we used three pOXA-48-carrying strains. These were a laboratory-adapted *E. coli* strain [β3914, a diaminopimelic acid (DAP) auxotrophic laboratory mutant of *E. coli* K-12], and two wild-type pOXA-48-carrying clinical strains recovered from hospitalized patients and belonging to STs responsible for between-patient plasmid dissemination in the same hospital (32): *K. pneumoniae* ST11, and *E. coli* ST10 (strains K93 and C165 described in references 32 and 36).

We first performed classical conjugation assays (one donor and one recipient strain in equal proportions) for all donor–recipient combinations, and measured conjugation frequencies as the ratio between transconjugants (TC) and total recipient cells (see Methods for details). Note that the donor *K. pneumoniae* ST11 could not be distinguished from three recipients of the same species, either by the phenotype in the differential medium or by antibiotic selection, and therefore conjugations could not be performed for these donor–recipient pairings. This analysis revealed a wide diversity of conjugation frequencies across recipient strains. Overall, *Klebsiella spp.* strains showed a significantly higher conjugation permissiveness than *E. coli* for all three donors (Fig. 2A, Kruskal-Wallis rank sum test: *E. coli* β3914, *X*^2^ = 45.8, *P* = 1.9 x 10^-11^; *E. coli* ST10, *X*^2^ = 23.8, *P* = 1.07 x 10^-6^; *K. pneumoniae* ST11, *X*^2^ = 6.98, *P* = 8.24 x 10^-3^). Moreover, two *E. coli* strains produced no TCs with any of the donors, whereas all *Klebsiella spp.* strains produced TCs in at least one of the conditions and were also the recipients with highest conjugation permissiveness for every donor. Finally, we also calculated the conjugation rates from the mating assays with *E. coli* ST10 as donor and the 20 recipient strains, using the end-point method (37) (Fig. S14). Results ruled out the possibility that differences in growth dynamics of donors, recipients and TCs could produce biases in the conjugation frequency results (37,38).

**Fig. 2.**
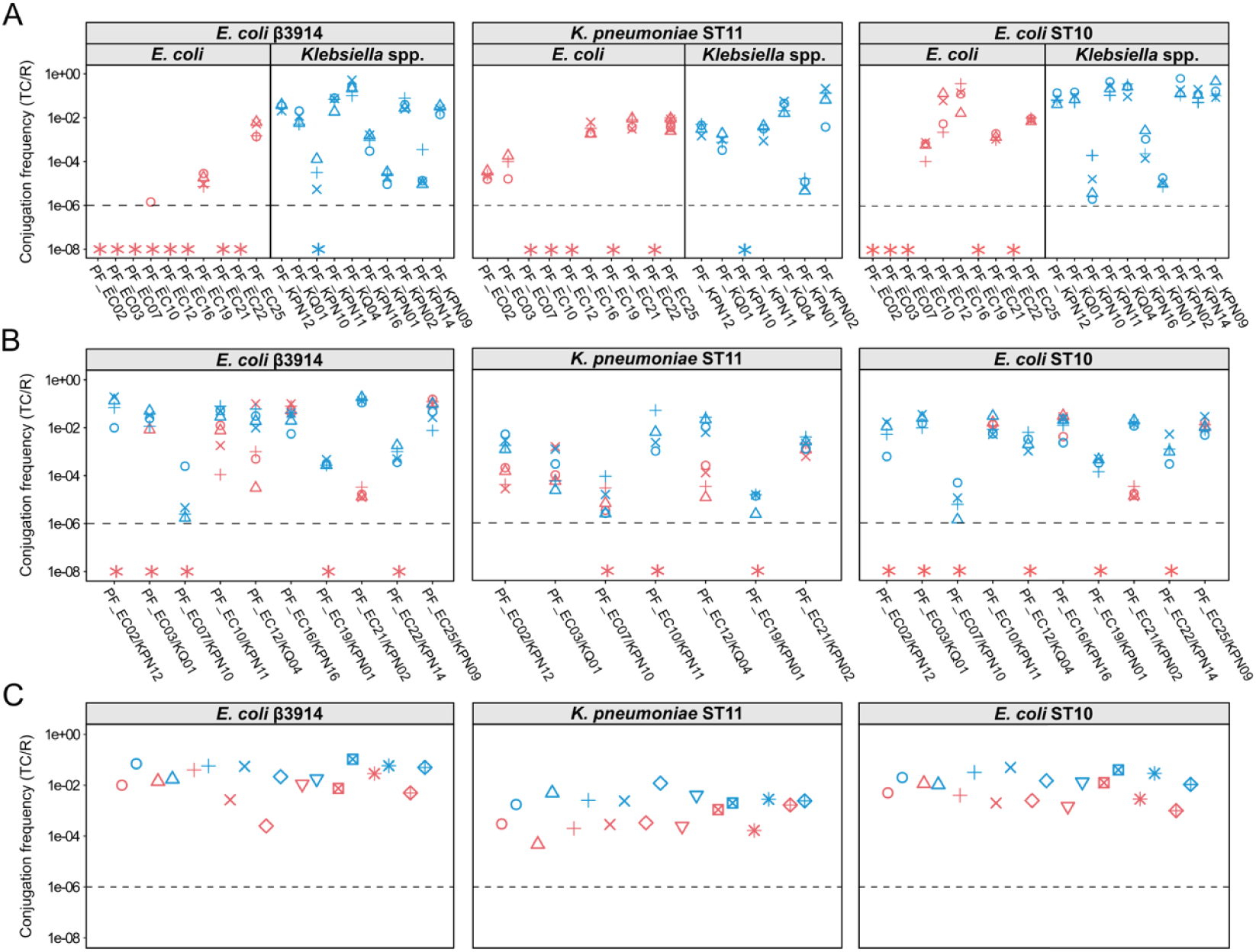
*In vitro* pOXA-48 conjugation dynamics in wild type enterobacteria. Conjugation frequencies (transconjugants/recipient) of a subset of 20 isolates from our clinical enterobacteria collection (10 *E. coli* and 10 *Klebsiella* spp., x-axis), obtained from three pOXA-48 donors (*E. coli* β3914, left; *K. pneumoniae* ST11, center; and *E. coli* ST10, right). (A) Conjugation frequencies obtained in conjugation assays with one donor and one recipient in equal proportions. Color represents the recipient species (red, *E. coli*; blue, *Klebsiella* spp.), symbols represent biological replicates (n = 4), and asterisks symbolize conjugation frequencies below the detection limit (dotted line). *Klebsiella spp.* exhibits higher conjugation frequencies than *E. coli* for the three donors. (Kruskal-Wallis rank sum test: *P* < 0.01). (B) Conjugation frequencies using one donor and pairs of *E. coli* and *Klebsiella* spp. as recipients. *Klebsiella spp.* is more conjugation-permissive than *E. coli*, regardless of the donor used (Wilcoxon signed-rank exact test: *P* < 0.01). (C) Conjugation frequencies obtained with a pool of recipients (one donor, 20 recipients). The results give a conjugation frequency value per species, and symbols represent biological replicates (n=9) whose values would correspond to the most successful strains of each species in plasmid uptake. *Klebsiella spp.* has higher conjugation frequencies than *E. coli*, regardless of the donor (Kruskal-Wallis rank sum test: *P* < 0.01).

The gut microbiota is a dense bacterial community that engages in complex ecological interactions. To analyze if community complexity could affect conjugation permissiveness in our clinical strains, we repeated the conjugation assays, but this time instead of using a single clone as the receptor, we used ten pairs of *E. coli*/*Klebsiella* spp. strains (Fig. 2B and Table S2). This analysis once again revealed great variability in reception frequencies and a higher overall conjugation permissiveness in *Klebsiella* spp. strains (Fig. 2B, paired Wilcoxon signed-rank exact test: *E. coli* β3914, *V* = 235, *P* = 0.017; *E. coli* ST10, *V* = 205, *P* = 0.01; *K. pneumoniae* ST11, *V* = 45, *P* = 1.2 x 10^-4^). In general, the conjugation frequencies observed agreed with those from the single conjugation assays, suggesting that ecological interactions did not markedly affect conjugation dynamics. However, one of the *E. coli* strains that produced no TCs as single receptor was able to acquire the plasmid (PF_EC07), which could be due to secondary conjugations from the *Klebsiella* spp. TC (39). Finally, to increase the complexity of the recipient community even further, we performed conjugation assays using a pool of all 20 strains as recipients (Fig. 2C). In this case, we could only distinguish between TCs at the species level in the differential medium. Once again, the results revealed *Klebsiella spp.* to be more conjugation-permissive than *E. coli*, regardless of the donor (Fig. 2C, Kruskal-Wallis rank sum test: *E. coli* β3914, *X*^2^ = 9.28, *P* = 2.32 x 10^-3^; *E. coli* ST10, *X*^2^ = 9.31, *P* = 2.35 x 10^-3^; *K. pneumoniae* ST11, *X*^2^ = 12.8, *P* = 3.49 x 10^-4^).

### Conjugation dynamics in a mouse model of gut colonization

To contrast our *in vitro* results in a more biologically relevant environment, we studied pOXA-48 conjugation dynamics in a mouse model of gut colonization (C57BL/6J). As donor, we used *K. pneumoniae* ST11 strain K93, since this clonal group is frequently responsible for gut microbiota colonization of patients in our hospital (32). As recipients, we used 6 strains representative of the variability in conjugation frequency observed *in vitro* (*E. coli* PF_EC10, PF_EC12, and PF_EC21; and *Klebsiella spp*. FP_KQ01, PF_KPN11, and PF_KPN12). Mice (n = 21) were treated orally with ampicillin (0.5 g/L), vancomycin (0.5 g/L), and neomycin (1 g/L) for one week to reduce colonization resistance, as we previously described (40), and the antibiotic treatment was stopped one day before inoculation. A total of 18 mice were inoculated orally with the donor and co-inoculated 2 hours later with the recipients (3 mice per recipient), and 3 mice were not inoculated with any bacteria, as controls. Fresh fecal samples were collected every 24 hours during 2 days. At the end of the second day, mice were sacrificed, and cecum samples were collected. Samples were processed and plated with antibiotic selection to determine colonization levels and the frequency of transconjugants, as described in Methods. Two independent assays were performed, two months apart, and the results of both assays are compiled below (Fig. 3 and source data).

**Fig. 3.**
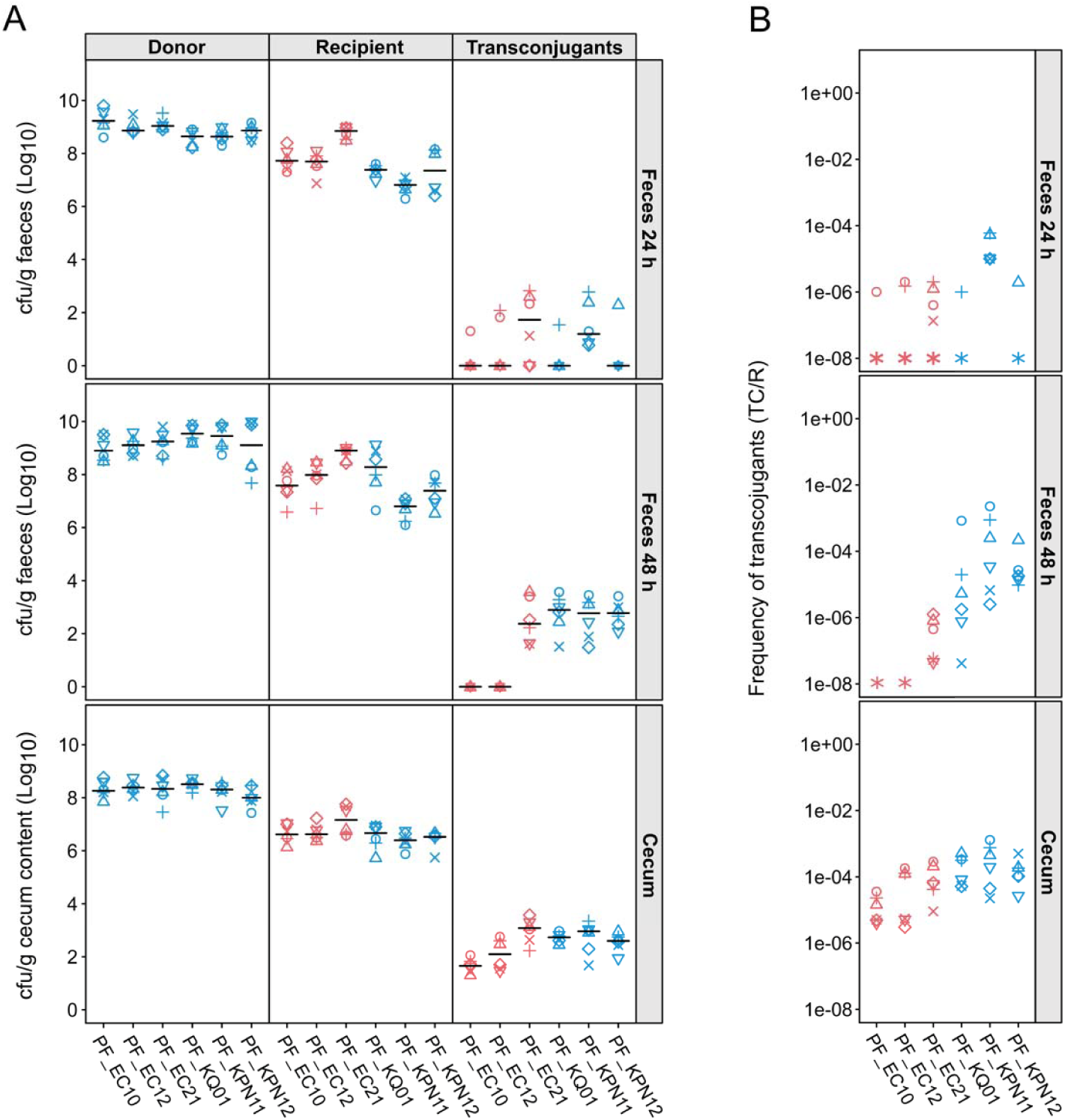
pOXA-48 conjugation dynamics in a mouse model of gut colonization. pOXA-48 transconjugant frequencies in mouse feces and cecum at 24 and 48 hours post colonization. (A) Colonization levels of the donor (*K. pneumoniae* ST11), recipients (3 *E. coli* and 3 *Klebsiella spp.* strains), and transconjugants, in feces at 24 and 48 hours post-inoculation (top and center) and in cecum content at 48 h (bottom). Species are indicated by colors (red, *E. coli*; blue, *Klebsiella spp.*) and biological replicates (n=6) by symbols. Black lines indicate the median values for the biological replicates. Donor and recipient colonization levels were high in both feces and cecum. (B) Transconjugant frequencies (transconjugants/recipients) in feces at 24 and 48 hours (top and center) and in cecum (bottom). Transconjugant frequencies were significantly higher in *Klebsiella* spp. than in *E. coli* in both feces and cecum at 48 hours (Kruskal-Wallis rank sum test in feces H = 24.6, *P* = 6.89 x 10^-7^; in cecum H = 10.2, *P* = 1.4 x 10^-3^).

Colonization levels were high in all inoculated mice at both 24 and 48 hours for donor and recipient strains (Fig. 3A), while no colonization was observed in the control mice. At 24 hours post-inoculation, the numbers of transconjugants were generally low, and no differences in transconjugant frequencies were detected between *E. coli* and *Klebsiella* spp. (Fig. 3B Kruskal-Wallis rank sum test H = 1.4, P = 0.265). At 48 hours post-inoculation, the levels of donors were similar in mice co-inoculated with *E. coli* or *Klebsiella* spp. recipient strains (Wilcoxon rank sum test: in feces, W = 2.71, P = 0.1; in cecum, W = 0.1, P = 0.752). However, despite the fact that the levels of *Klebsiella spp.* recipients were slightly lower than those of *E. coli* recipients (Wilcoxon rank sum test: in feces, W = 245, P = 0.008; in cecum, W = 4.9, P = 0.026), *Klebsiella spp.* showed significantly higher frequencies of transconjugants (Fig. 3B, Kruskal-Wallis rank sum test: in feces, *H* = 24.6, *P* = 6.89 x 10^-7^; in cecum, *H* = 10.2, *P* = 1.4 x 10^-3^). In summary, the *in vivo* results correlated qualitatively with those from the *in vitro* conjugation assays (Fig. 2A and 3B), and *Klebsiella spp.* strains showed higher levels of transconjugants than *E. coli* strains on average.

### Simulating plasmid-host associations in complex populations

We previously developed a mathematical model to study how pOXA-48 fitness effects impact plasmid maintenance in complex bacterial communities (34). That model allowed us to predict the competitive fitness of each strain, with and without pOXA-48, based on growth-kinetic parameters that were calibrated from experimental growth curves obtained from single clones. In the present study, we extended this model to include the new variables informed by the experimental results: conjugation permissiveness and resistance levels (Fig. 4A,B). Specifically, we used the conjugation rates values obtained using *E. coli* ST10 as donor, and resistance levels to AMC (see Supporting Information for a complete description of the model and its parametrization). This computational model allowed us to track the distribution of plasmids in response to selection at species and strain resolution.

**Fig. 4.**
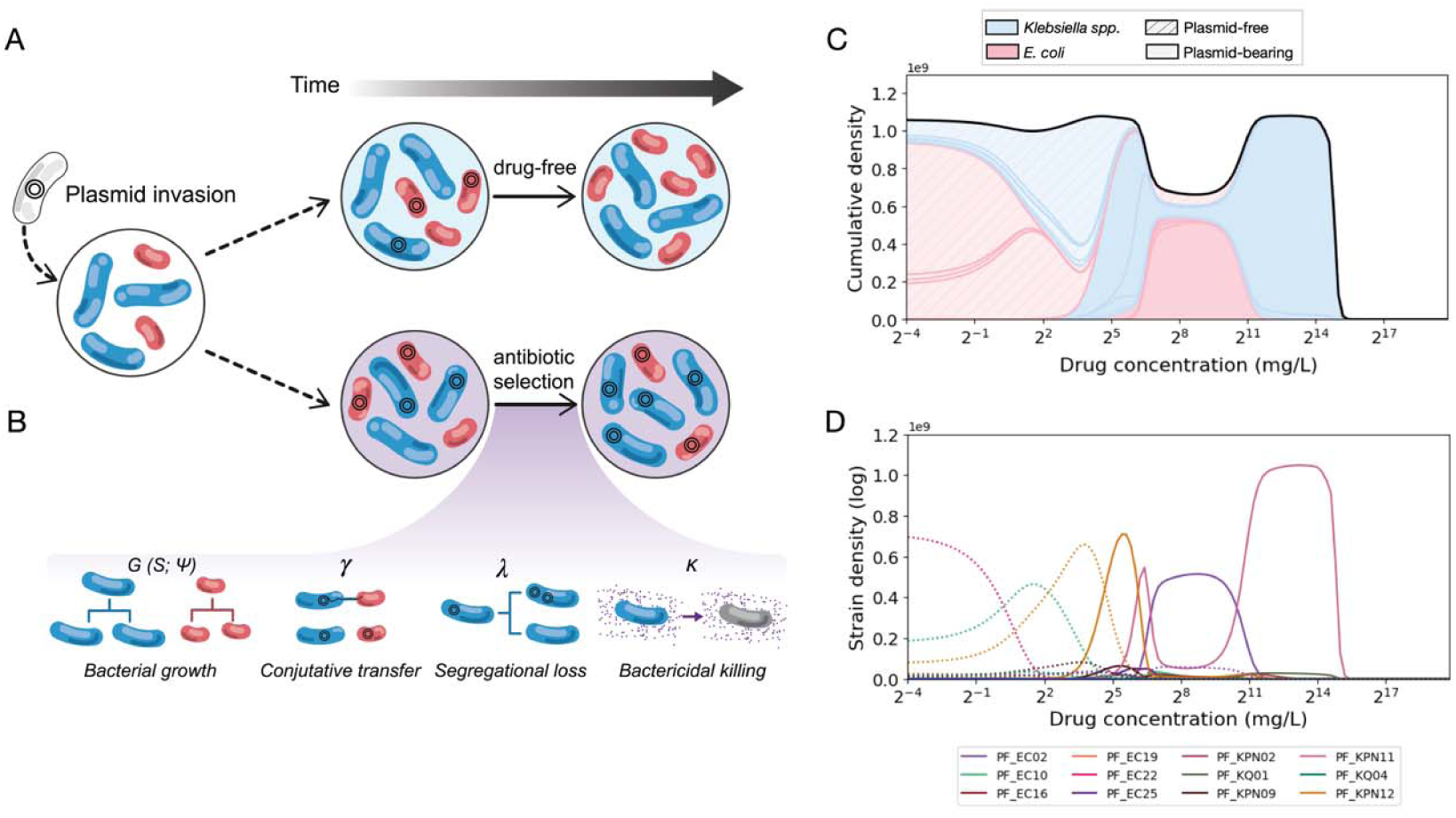
Computational model of plasmid population dynamics. (A) Diagram illustrating the computational approach employed to assess the distribution of plasmids within mixed bacterial populations under a range of selective pressures (see Supporting Information). (B) The model simulates various ecological processes that influence plasmid dynamics, enabling the exploration of their behavior under different conditions, including bacterial growth, modeled as a Monod term G (S; ψ), conjugative transfer occurring at a rate Y, segregational loss occurring at a rate λ, and antibiotic bactericidal killing k (Table S1). (C) Population dynamics simulated across a range of drug concentrations, starting with an initial population of 10 *E. coli* and 10 *Klebsiella* spp. strains. Predominantly, plasmid-free bacteria emerge at low drug levels, while plasmid-bearing *Klebsiella* spp. prevail under high drug concentrations. (D) Strain-level resolution in the simulations of the multistrain dose-response. In conditions of low drug concentrations, the plasmid-free strains PF_EC22 (ST131) and PF_KPN12 (ST307) are predominant. In high drug concentration environments, the strain PF_KPN11 (ST307) appears to contribute to the preservation of plasmid stability in these communities.

We first simulated simple pairwise competition experiments between the 10 *Klebsiella* spp. and 10 *E. coli* strains in our collection with experimentally determined conjugation rates and AMR levels (Table S1). Each competition experiment started with a 1:1 ratio, and a 50% plasmid frequency in each strain (see Supporting Information). To assess the role of antibiotics in the distribution of plasmids in the population, we performed all pairwise competition experiments for a range of selective pressures (Fig. S5-S7). At low and intermediate antibiotic concentrations, the plasmid was preferentially associated with *E. coli*. In contrast, as the selective pressure increased, the proportion of plasmid-carrying *Klebsiella* spp. strains within the pairs correspondingly increased, reflecting the relatively lower resistance levels exhibited by *E. coli* strains (Fig. S5-S7).

We next used the model to analyze the impact of AMR levels and conjugation permissiveness in pOXA-48 distribution in complex communities, such as those in the gut microbiota of hospitalized patients (Fig. 4 and S8). Our previous epidemiological studies revealed that patients are commonly colonized by pOXA-48-carrying nosocomial enterobacterial clones (32). During colonization, the plasmid spreads horizontally to resident enterobacteria in the gut, and different clones can subsequently maintain the plasmid in the community, producing long term pOXA-48 gut carriage (32). To simulate the invasion and dissemination within the gut microbiota community, we used our mathematical model to initialize multistrain simulations with only one member of the community carrying pOXA-48 (starting at a frequency of 0.01% of the total density, Fig. 4A,B). We conducted 10-day simulated serial dilution experiments with a starting community including the 20 strains (the 10 *E. coli* and 10 *Klebsiella spp.*). We then explored the effects of various antibiotic concentrations on community compositions (Fig S8), aiming to evaluate the plasmid dynamics and to identify the successful associations between plasmid and hosts. At low drug concentrations, we found that, as the selective pressure increases, the population shifts from being dominated by plasmid-free *E. coli* to contain mostly plasmid-free *Klebsiella* spp. (Fig. 4C). At intermediate and high drug concentrations, plasmid-carrying strains dominate, with only *Klebsiella* spp. strains being able to persist at very high concentrations. At the strain-level, the simulations revealed that the species-level dynamics were driven by a small subset of successful strains, which played a key role in stabilizing the plasmid in the communities (Figs 4D). At intermediate and high drug concentrations, plasmid-carrying PF_KPN12 (ST307), PF_EC02 (ST453), and PF_KPN11 (ST307) were the most prevalent strains. At very high antibiotic concentrations pOXA-48-carrying PF_KPN11 (ST307) clearly dominated the population.

Finally, we followed a different approach to further evaluate if specific bacterial strains played an important role in maintaining plasmid stability within the community. We used an iterative knock-out simulation approach where strains were sequentially and randomly removed from the 20-strain community, with an evaluation of plasmid stability after each removal. This approach revealed the important role of strains PF_EC03 (ST131), PF_EC16 (ST38), PF_KPN11 (ST307) in antibiotic-free environments (Fig. S9). In high-antibiotic-concentration environments, PF_KPN02 (ST485), PF_KQ01, and again, PF_KPN11 (ST307) played key roles in maintaining plasmid stability (Fig S9).

Taken together, our results underscore the critical role of successful plasmid-host associations in securing plasmid stability in mixed microbial populations. Specifically, we observed that certain strains in our pool, such as PF_KPN11 (ST307), are crucial for the dissemination and maintenance of plasmid pOXA-48 in the communities.

### *Klebsiella*-derived capsules are associated with plasmid carriage in *E. coli*

The conjugation experiments showed a higher permissiveness for pOXA-48 acquisition in *Klebsiella* spp. than in *E. coli*, as well as high variability between strains of the same species. We sought to investigate the genetic traits that could explain these differences. We analyzed the genomes of the 20 recipient strains for an array of traits known to affect conjugation efficiency: CRISPR arrays, restriction-modification (RM) systems and other phage defense systems that could target pOXA-48, plasmid incompatibilities, and type VI secretion systems (T6SS, known to mediate bacterial competition, which may affect conjugation frequencies). However, none of these traits accounted for the observed differences in pOXA-48 acquisition (Fig. S15, Table S3).

Capsules have been reported to hinder DNA transfer and possibly constitute a barrier to plasmid acquisition (41, 42). We used two software tools to search for capsular systems: Kaptive, to detect and type *Klebsiella* capsular loci, and CapsuleFinder, to identify other capsular systems. With the single exception of the non-capsulated PF_KPN14 strain, each recipient strain encoded a specific set or type of complete capsular loci (Table S3). To assess whether similar sets of capsular loci were associated with higher or lower conjugation permissiveness, we calculated a weighted gene repertoire relatedness (wGRR) score. Although we found no clear pattern for the *Klebsiella* spp. recipients, the *E. coli* strain with the highest overall pOXA-48 reception permissiveness across the three different donors, PF_EC25, encoded a *Klebsiella* capsular locus (KL35, Fig. 5A, Table S3).

**Fig. 5.**
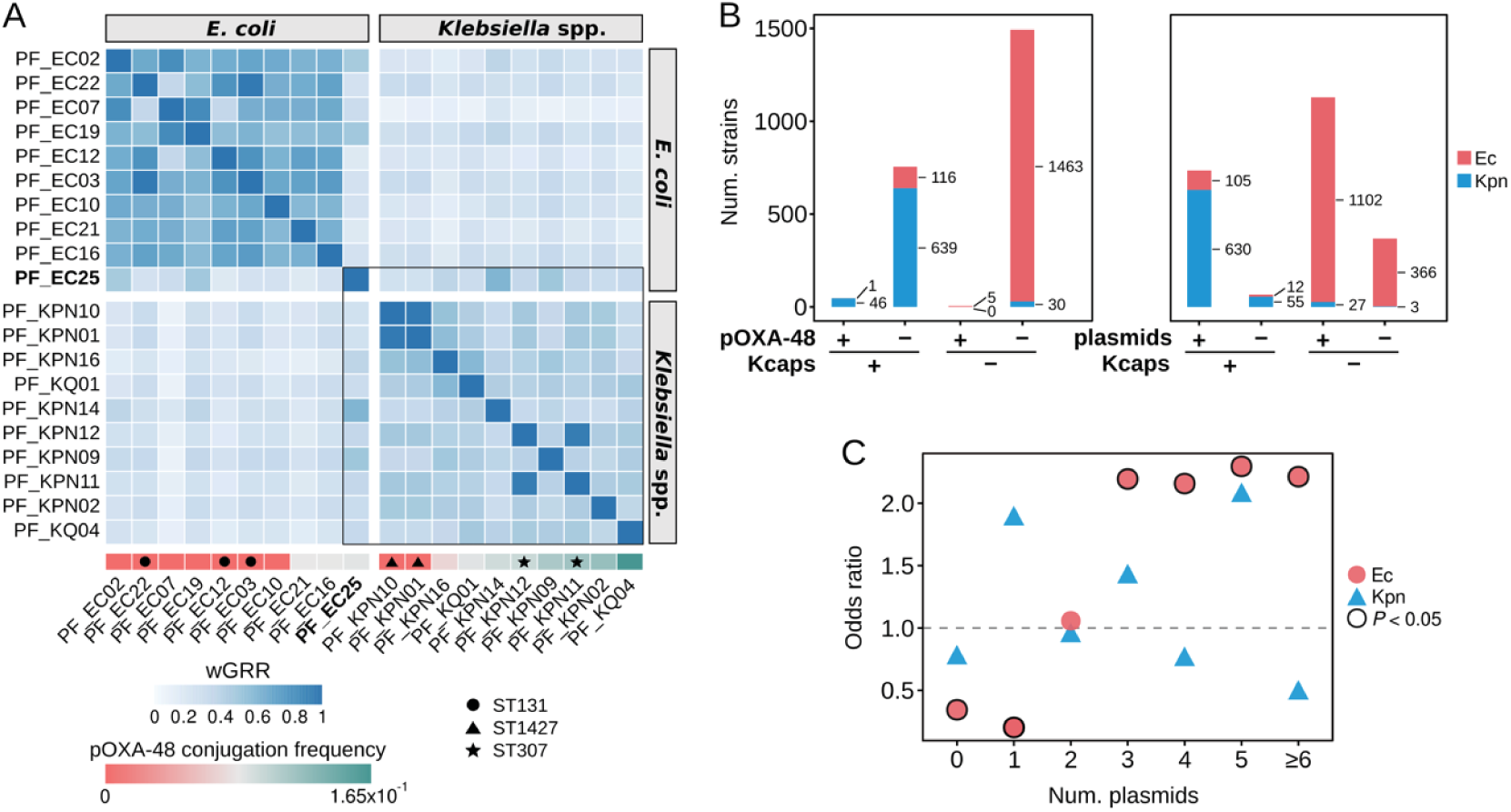
Associations between capsular systems and plasmid carriage. Analysis of capsular systems and plasmid prevalence in our recipient strains and in *E. coli* and *K. pneumoniae* genomes obtained from RefSeq. (A) Heatmap of weighted gene repertoire relatedness (wGRR) scores between capsular sets (*Klebsiella*-derived capsules and/or other capsular systems) encoded by the recipient strains used in conjugation experiments. A wGRR score of 1 means that all proteins from the capsular set encoding the fewest number of proteins have a homolog in the capsular set it is compared with. A score of 0 indicates there are no homologs between the capsular sets of the two compared strains. Recipients are ordered by median pOXA-48 conjugation frequency across replicates and donors. The PF_EC25 strain, highlighted in bold, has the highest median conjugation frequency of any *E. coli* strain and is the only *E. coli* strain encoding a *Klebsiella*-derived capsule. Strains from the same sequence type (ST) are marked with a symbol. The black square indicates strains with a *Klebsiella*-derived capsule. (B) Number of *E. coli* (Ec) and *K. pneumoniae* (Kpn) strains from the RefSeq database analyzed for the association between the presence of *Klebsiella*-derived capsules (Kcaps) and carriage of pOXA-48 (left) or plasmids in general (including pOXA-48; right). In total, the RefSeq database includes 1,585 Ec and 715 Kpn strains. (C) Association between the presence of a genome-encoded *Klebsiella*-derived capsule and the number of plasmids carried in the *E. coli* (Ec) and *K. pneumoniae* (Kpn) RefSeq strains. The chart shows odds ratio (Fisher’s exact test) of plasmid carriage for strains encoding a *Klebsiella*-derived capsule *vs* strains encoding other capsular systems (not from *Klebsiella*). Values above 1 (horizontal dashed line) indicate a positive association, and values below 1 a negative association, between encoding *Klebsiella*-derived capsules and carrying the indicated number of plasmids. In all cases, the number of strains carrying each number of plasmids is >100. A significant Fisher’s test (*P* < 0.05) is indicated by a black outline.

To determine if *Klebsiella*-derived capsules better facilitate pOXA-48 acquisition than other capsular systems, we analyzed their association in the complete genomes of 1,585 *E. coli* and 715 *K. pneumoniae* strains (RefSeq), of which 6 *E. coli* strains and 46 *K. pneumoniae* strains carry pOXA-48 (Fig. 5B, Table S4). We used phylogenetic logistic regression (*phyloglm*) to account for phylogenetic dependency of the significant associations (*P* < 0.05, Fisher’s exact test; Table S6). The presence of *Klebsiella*-derived capsules was associated with pOXA-48 carriage only when *E. coli* and *K. pneumoniae* were analyzed together (Table S6, *phyloglm P* = 0.0016). We next investigated the relationship between the presence of *Klebsiella*-derived capsules and plasmid carriage more generally, finding significant associations when analyzing *E. coli* and *K. pneumoniae* together (Table S6, *phyloglm P* < 2 x 10^-16^) and when analyzing *E. coli* separately (Table S6, *phyloglm P* = 2.2 x 10^-5^). Moreover, *E. coli* strains encoding other capsular systems (not derived from *Klebsiella*) were more likely to be plasmid-free or to carry only one plasmid, whereas *E. coli* encoding *Klebsiella*-derived capsules had an increased likelihood of carrying multiple plasmids (Fig. 5C). These results suggest that although capsules generally obstruct conjugation (41, 42), certain types of *Klebsiella*-derived capsules could be more permissive than other capsule types to plasmid acquisition by *E. coli*.

## Discussion

In this work, we performed a comprehensive analysis of pOXA-48 effects in clinical enterobacteria. One of the main strengths of this study is the relevance of the experimental system. We used an epidemic carbapenem resistance plasmid, pOXA-48, and a collection of wild-type, multidrug resistant, enterobacterial isolates recovered from hospitalized patients. Using this system, we analyzed pOXA-48-associated AMR levels and conjugation permissiveness both *in vitro* and *in vivo* (Figs. 1-3). The key advantage of experimentally determining plasmid-associated phenotypes in a collection of wild type clinical enterobacteria is that we can confidently feed these variables into mathematical models. We developed a model including three key variables informed by our experimental results: bacterial growth dynamics with and without pOXA-48 (34), conjugation permissiveness (Fig. 2 and S14), and AMR levels (Fig. 1). We used this model to investigate pOXA-48 dynamics in complex bacterial communities under different levels of selection, aiming to simulate pOXA-48 invasion of a gut microbiota (Fig. 4).

The critical result of our study is that pOXA-48-associated phenotypes vary significantly across different wild type bacterial hosts. We had previously shown that pOXA-48 produces variable fitness effects in the absence of antibiotics (34, 43), and in this new study we reveal similar variability in pOXA-48-associated AMR levels and conjugation permissiveness. This variability is especially evident for horizontal transmission, with conjugation frequencies differing by orders of magnitude across different recipients. Our simulations revealed that the variability of plasmid-associated phenotypes dramatically impact plasmid distribution in polyclonal bacterial communities. Specifically, a single successful pOXA-48-bacterium association is sufficient to allow for plasmid invasion and stable maintenance in a complex population for a wide range of antibiotic concentrations (Fig. S8). These jackpot associations, combining high growth rates, high AMR levels, and high conjugation permissiveness, can act as pOXA-48 super-sinks in the gut microbiota of patients, maintaining and further disseminating the plasmid. Interestingly, these successful pOXA-48-carrying clones are likely to produce long-term colonization of the gut microbiota of patients, as we have recently reported (32). Long-term colonization may in turn lead to the dissemination of carbapenemases from clinical settings towards the community, which represents a critical step in AMR epidemiology.

One of the main goals of this study was to understand the bias in pOXA-48 distribution across enterobacteria. Despite having a broad host range and producing similar fitness effects in *E. coli* and *Klebsiella* spp. (34, 43), pOXA-48 is usually associated with specific *K. pneumoniae* clones in hospitals (32, 33). Our results help to explain this bias. We experimentally showed that, compared with *E. coli*, *Klebsiella* spp. clones generally have a higher AMR level associated with pOXA-48 carriage and a greater ability to take up pOXA-48 by conjugation. Crucially, our simulations of both pairwise competitions and complex bacterial communities revealed that pOXA-48 is mostly associated with *Klebsiella* spp. under high antibiotic pressure. Taken together, these results help to explain why pOXA-48 is usually associated with *K. pneumoniae* in the gut microbiota of hospitalized patients, which is a complex community frequently exposed to high antibiotic pressure. At the strain level, our model also provides interesting information. The most relevant result emerging from our simulations is that *K. pneumoniae* strains belonging to ST307 (PF_KPN12 and specially PF_KPN11) are responsible for pOXA-48 maintenance in the bacterial communities. Importantly, ST307 is a clonal group that is emerging as major cause of nosocomial outbreaks in many different regions around the world, and is usually associated with plasmid-mediated extended-spectrum ß-lactamases (ESBLs) and carbapenemases (33, 44, 45). In Spain, ST307 is actually becoming the most concerning pOXA-48-carrying clonal group in hospitals, replacing other high-risk *K. pneumoniae* STs, such as ST11 (32, 46). Therefore, our simulations based on data obtained from clinical strains which were originally pOXA-48-free were able to mirror the real dynamics observed in our hospital. These results suggest that the experimentally determined phenotypes: fitness with and without plasmid, AMR levels and conjugation permissiveness, could help to predict the evolution of plasmid-mediated AMR in clinical settings.

Given the importance of the variability of plasmid-associated phenotypes in the evolution of plasmid-mediated AMR, a critical research direction in the field will be to characterize the molecular basis underlying these differences. In this study, we showed that the presence of *Klebsiella*-derived capsules, compared to other capsular systems, is associated with a higher plasmid frequency in an *E. coli* clone (Fig. 5). We also showed that pOXA-48 PCN is higher in *Klebsiella* spp. than in *E. coli* clones, probably contributing to the higher AMR level observed in pOXA-48-carrying *Klebsiella* spp. strains (Fig. 1). Further work will be needed to characterize the molecular basis and significance of these and other specific interactions. Understanding the plasmid and host factors that determine this variability will help us not only to predict, but also hopefully to counteract successful associations between AMR plasmids and high-risk bacterial clones.

## Methods

### pOXA-48_K8 plasmid, strains, and growth conditions

We used 50 wild type pOXA-48-free enterobacterial strains recovered from the gut microbiota of hospitalized patients (Table S2, partially overlapping the collection in (34)). These strains belonged to the R-GNOSIS collection, which was constructed in our hospital during an active surveillance-screening programme for ESBL/carbapenemase-producing enterobacteria (from March 4th, 2014 to March 31st, 2016, R-GNOSIS-FP7-HEALTH-F3-2011-282512, https://cordis.europa.eu/project/id/282512/reporting/es, approved by the Ramón y Cajal University Hospital Ethics Committee, Reference 251/13). These strains are representative of *E. coli* and *Klebsiella* spp. diversity in the R-GNOSIS collection, and we define them as ecologically compatible with plasmid pOXA-48, because they were obtained from patients on hospital wards where pOXA-48-carrying enterobacteria were commonly isolated during the same period. To analyze the pOXA-48-associated AMR level, we selected 33 isogenic strains pairs (15 *E. coli* and 18 *Klebsiella* spp.) with (TC) or without pOXA-48 plasmid (PF) from the collection. Plasmid-bearing strains carry the most common pOXA-48-like plasmid variant from our hospital, pOXA-48_K8 (32). For the conjugation frequencies determination, we selected, as recipients, 10 *E. coli* and 10 *Klebsiella spp.* pOXA-48-free isolates that cover the phylogenetic diversity of the collection (Fig. S10) (32) and showed similar β-lactam resistance levels. As plasmid donors we selected three pOXA-48_K8-carrying strains: *E. coli* β3914 (47), *K. pneumoniae* K93, and *E. coli* C165 (32, 36). Bacterial strains were cultured in LB at 37 °C with continuous shaking (250 r.p.m), and on LB or HiCrome^TM^ UTI (Himedia Laboratories, India) agar plates at 37 °C.

### Determination of minimum inhibitory concentration (MIC)

We determined the MIC of AMC (Normon, Spain), ERT (Merck Sharp & Dohme B.V., Netherlands), IMP (Fresenius Kabi, Germany) and MER (SunPharma, India) in all plasmid-carrying and plasmid-free clones selected for this study following a modified version of the agar dilution protocol (48, 49). This method allows us to test a large number of isolates simultaneously under identical conditions and to test a wide range of AMC concentrations, since this β-lactam antibiotic degrades fast in liquid medium (50). We prepared pre-cultures of plasmid-free and plasmid-carrying strains by inoculating single independent colonies into LB broth in 96-well plates and overnight incubation at 37 °C with continuous shaking (250 r.p.m.). We spotted 10 µl of each overnight culture onto LB agar plates with increasing concentrations of AMC (from 4 mg/L to 32,768 mg/L), ERT (from 0,25 mg/L to 512 mg/L), IMP (from 0.25 mg/L to 1,024 mg/L) or MER (from 0.25 mg/L to 32 mg/L), and incubated them overnight at 37 °C. We determined the MIC of each strain as the lower antimicrobial concentration able to inhibit the growth of each strain. We performed three biological replicates per MIC determination in three independent days, and we used the median value of the three replicates as the MIC.

### *In vitro* determination of conjugation frequencies and rates

To determine the *in vitro* conjugation frequencies, we selected, as recipient strains, 10 *E. coli* and 10 *Klebsiella spp.* pOXA-48-free isolates, from our collection of ecologically compatible enterobacterial isolates (Fig. S10). As plasmid donors we used three pOXA-48_K8-carrying strains: *E. coli* β3914, a counter-selectable diaminopimelic acid auxotrophic laboratory strain (47); *K. pneumoniae* K93, a pOXA-48-carrying ST11 strain; and *E. coli* C165, a pOXA-48-carrying ST10 strain (32, 36). To counter-select the two natural donors, we transformed the recipients with pBGC plasmid, a non-mobilizable *gfp*-carrying small plasmid that encodes a chloramphenicol resistance gene (34). It was not possible to introduce pBGC into three *Klebsiella spp.* isolates (PF_KPN09, PF_KPN14 y PF_KPN16), due to their level of constitutive resistance to chloramphenicol. We streaked donor and recipient strains from freezer stocks onto LB agar plates with or without ertapenem 0.5 mg/L –and 0.3 mM DAP for *E. coli* β3914/pOXA-48_K8–, respectively, and incubated at 37 °C overnight. We prepared pre-cultures of donors and recipients by inoculating single independent colonies in 15-ml culture tubes containing 2 ml of LB broth and overnight incubation at 37 °C with continuous shaking (250 r.p.m.). To perform classic conjugation assays, we mixed one donor and one recipient at equal proportions (each donor with each recipient, independently, with a total of 4 recipient biological replicates by donor), plated the mixture as a drop onto LB agar plates – supplemented with DAP 0.3mM for *E. coli* β3914/pOXA-48_K8 donor– and incubated them at 37 °C during only 4h. A short mating experiment (4 hours) reduces the chances of secondary conjugation events from transconjugants to recipients, but we cannot rule out this possibility. Also, 4 hours mating reduces the impact of potential differences in donor, recipient, and transconjugant growth rates on conjugation frequency determination. After incubation, we collected the biomass, resuspended it in 1 ml of NaCl 0.9 % and performed 1/10 serial dilutions up to 10^-6^. We estimated the final densities of transconjugants and recipients by plating 10 µl of each dilution on HiCrome^TM^ UTI agar plates (HIMEDIA laboratories, India) with or without AMC 256 mg/L, respectively, and with chloramphenicol 50 mg/L. We plated the dilutions as drops, and each drop was subsequently allowed to drain down the plate, this way we can separate the colonies so they are easily distinguishable and quantifiable. We performed the donor counter-selection by the absence of DAP in the medium for *E. coli* β3914/pOXA-48_K8, or by selection with chloramphenicol for clinical *K. pneumoniae* ST11 and *E. coli* ST10. Finally, we calculated the conjugation frequencies per recipient following the formula below:

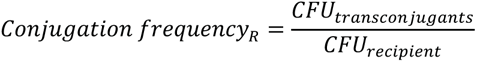

where *CFU_transconjugants_* are the transconjugant colony-forming units obtained from the number of colonies in the plate (corrected by the dilution factor); and *CFU_recipient_* are the number of recipient colony-forming units minus transconjugants. Note that in the cases where no TC colonies were detected in the plate, we calculated a threshold conjugation frequency assuming the presence of a single colony in the total cell suspension, thus establishing a detection limit of 10^-6^. Conjugation rates were calculated for the experiments performed using *E. coli* ST10 as a donor and the 20 recipient strains. For these mating experiments, the number of donors, recipients and transconjugants were determined both before and after the mating assays as described above, and the conjugation rates were determined using the end-point method for solid surfaces (37).

To perform conjugation assays with recipient pairs we randomly grouped the recipients in pairs so that each pair comprised one *E. coli* and one *Klebsiella spp.* recipient. We used a modified version of the above protocol, in which we mixed recipient pre-cultures of each pair at equal proportions and we subsequently mixed these recipient mixtures with the pre-culture of the donor at the same proportion, forming the final conjugation reaction (each recipient pair with one donor per time, with 4 biological replicates of recipient pairs). We determined the recipients and transconjugants densities by antibiotic selection with AMC 256 mg/L, (to select transconjugants vs. recipients) and chloramphenicol 50 mg/L (to counter-select donors), and distinguished between recipients using the differential chromogenic medium HiCrome^TM^ UTI agar. We calculated the conjugation frequency per each receptor using the formula above.

For conjugation assays with a pool of all the recipients, we mixed the same pre-culture volume of all recipient strains with one donor per experiment, following the same protocol previously described. Nine biological replicates were performed.

### *In vivo* conjugation

For the determination of conjugation frequencies of clinical isolates *in vivo*, we performed conjugation assays using a mouse model of gut colonization. We selected a subset of 6 pBGC-carrying recipient strains from the *in vitro* conjugation assays: 3 *E. coli* (PF_EC10, PF_EC12 and PF_EC21) and 3 *Klebsiella spp.* (PF_KQ01, PF_KPN11 and PF_KPN12) strains. As plasmid donor we selected the natural pOXA-48_K8-carrying isolate, *K. pneumoniae* ST11, K93 (32). We carried out the experiment described below twice, 2 months apart and under identical conditions.

#### Mouse model and housing conditions

We used 6-week-old C57BL/6J female mice (n = 21) purchased from Charles River Laboratories and housed them with autoclave-sterilized food (a 1:1 mixture of 2014S Teklad Global diet and 2019S Teklad Global Extrused 19% Protein Rodent Diet from Envigo) and autoclave-sterilized water. Temperature was kept at 21 ± 2 °C and humidity was maintained at 60–70%, in 12 h light/dark cycles. To allow intestinal colonization of the donor and recipient strains, we disrupted the intestinal microbiota by administering ampicillin (500 mg/L), vancomycin (500 mg/L) and neomycin (1 g/L) in the drinking water for one week, as previously described (40). Water with antibiotics was changed every 3-4 days to avoid reduction of antibiotic activity. During antibiotic treatment, mice were co-housed (3-6 mice per cage). After 1 week of treatment, antibiotics were removed from the drinking water and mice were individually housed to avoid bacterial transfer between mice due to coprophagia during the experiment.

#### Mice inoculation with donor and recipient strains

1 day after antibiotic treatment removal, 18 out of 21 mice were inoculated by oral gavage with 10^7^ CFU of the donor strain, *K. pneumoniae* ST11. Two hours later, each donor-inoculated mouse was co-inoculated by oral gavage with 10^7^ CFU of a recipient strain (3 mice per recipient). This way, from the initial 21 mice, 18 of them were inoculated with donor and recipient bacteria (6 different recipients, 3 mice per recipient) and 3 mice remained uninoculated as control.

#### Sample collection and bacterial levels determination

After antibiotic treatment and prior to inoculation, we collected fresh fecal samples from each mouse as to control for the absence of bacteria able to grow on the selective medium used for our bacteria of interest. We weighted the feces and resuspended them in 1 ml PBS until a homogeneous mixture was obtained. Then, we diluted mixtures 1:100,000 and plated 100 µl on HiCrome_TM_ UTI agar plates with ampicillin (400 mg/L) and vancomycin (1 mg/L). We incubated the plates overnight at 37 °C. After incubation, bacterial growth was found to be absent. During the assay, we collected fresh fecal samples from each mouse 24 and 48 h post inoculation and processed samples as described before. To determine the donor, recipients and transconjugants densities, we plated 100 µl of different dilutions onto HiCrome_TM_ UTI agar plates with: i) vancomycin (1 mg/L) and AMC (256 mg/L) to detect the donor; ii) vancomycin (1 mg/L) and chloramphenicol (50 mg/L) to detect recipients; and iii) vancomycin (1 mg/L), AMC (256 mg/L) and chloramphenicol (50 mg/L) to determine the presence of transconjugants. We calculated the colonization levels of the members of each conjugation assay with the following formula:

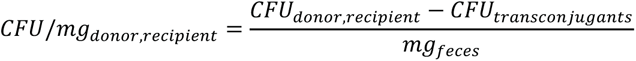

From the levels of recipients and transconjugants we calculated the conjugation frequencies per recipient as described before in the *in vitro* protocol. At 48 h post inoculation, after fresh fecal samples collection, we sacrificed mice and obtained samples of cecum content by manual extrusion. We performed sample processing, bacterial levels determination and conjugation frequencies determination as described above.

### DNA extraction and genome sequencing

We isolated the genomic DNA of the strains used in this work using the Wizard genomic DNA purification kit (Promega, WI, USA) and quantified using the Qubit 4 fluorometer (ThermoFisher Scientific, MA, USA). Whole genome sequencing was performed at the Wellcome Trust Centre for Human Genetics (Oxford, UK) using the Illumina HiSeq4000 or NovaSeq6000 platforms with 125 or 150 base pair (bp) paired-end reads, respectively. Read data is available under the BioProject PRJNA803387.

### Processing of sequencing data

All Illumina reads were trimmed with a quality threshold of 20 and reads shorter than 50 bp and adapters were removed using Trim Galore v0.6.6 (https://github.com/FelixKrueger/TrimGalore). PF and TC genomes were *de novo* assembled with SPAdes v3.15.2 (51) in--isolate mode and with the--cov-cutoff flag set to auto. Assembly quality was assessed with QUAST v5.0.2 (52). TC were confirmed to correspond to their PF strain by checking mutations and plasmid content with Snippy v4.6.0 (https://github.com/tseemann/snippy) and ABRicate v1.0.1 (PlasmidFinder database, https://github.com/tseemann/abricate), respectively. Snippy was also used to confirm isogenicity of the acquired pOXA-48 plasmids. Multilocus sequence typing was performed with MLST v2.21.0 (https://github.com/tseemann/mlst).

### Construction of phylogenetic trees

Mash distance phylogenies were constructed for the 25 *E. coli* and 25 *Klebsiella* spp. PF strains from their whole-genome assemblies using mashtree v1.2.0 (53) with a bootstrap of 100 (Fig. S10).

### Plasmid copy number (PCN) estimation

PCN of pOXA-48 was determined for the 33 TC strains and the 200 wild-type *E. coli* and *Klebsiella* spp. R-GNOSIS strains (BioProject PRJNA626430) carrying pOXA-48 variants sharing 72% of the pOXA-48_K8 core genome (32, 36). PCN was estimated from read coverage data as the ratio of plasmid median coverage by chromosome median coverage (54, 55). Since the genome assemblies are fragmented, the median coverage of the chromosome was calculated from the first three, largest contigs (total size sum of first three contigs 0.3-4.0 Mb), which were confirmed to correspond to chromosomal sequences by using BLASTn against the NCBI nr nucleotide database. The median coverage of pOXA-48 was calculated from the contig containing the IncL replicon, as identified with ABRicate v1.0.1 with the PlasmidFinder database, which generally corresponds to the largest pOXA-48 contig (size 11.0-59.8 Kb). For computing the PCN, the Illumina trimmed reads were first mapped to their respective genome assembly using BWA MEM v0.7.17 (56). Strain K25 from the wild-type pOXA-48-carrying collection was removed from the analysis due to truncated reads. Samtools depth v1.12 (57) with the-a flag was used to obtain read depths at each genomic position and the median read coverage for pOXA-48 and chromosome was computed with GNU datamash v1.4 (gnu.org/software/datamash).

### Comparative genomic analysis of recipient strains

We analyzed the genomes of the recipient strains to find traits that could explain the observed differences in pOXA-48 acquisition. First, the draft genomes of the 20 recipients and three donors (*E. coli* ST10 C165 and *K. pneumoniae* ST11 K93 from Bioproject PRJNA626430; for *E. coli* β3914, the sequence of the ancestral K-12 strain, NC_000913.3) were annotated with Prokka v1.14.6 (58) with default settings. A summary of the results can be found in Table S3.

Restriction-modification (RM) systems were searched using two approaches. First, the protein sequences of the recipient and donor strains were blasted against the protein “gold standard” REBASE database (downloaded February 2022, (59)) using BLASTp v2.9.0 (60). The results were filtered with a criteria based on (61). Briefly, hits with type I and IIG RM systems were kept if the percentage of identity and alignment was 100%, since it was observed that recognition sites varied between enzymes with only a few mismatches. For type II systems, a threshold of 70% identity and 90% alignment was used, as it was previously reported that these systems with identity over 55% generally share the same target motifs. For type III and type IV this threshold was higher, as reported in the same study, and was set at 90% identity and 90% alignment. After filtering, some proteins had hits with more than one enzyme. For these cases, the best hit or the hit with the enzyme of the corresponding organism (*E. coli* or *Klebsiella* spp.) was kept. The DefenseFinder webserver (accessed February 2022, (62, 63)) was also used to search for RM systems in the proteomes of recipients and donors. Then, the REBASE hits and the RM systems detected by DefenseFinder were unified by protein ID. Enzymes or RM systems present in all strains were discarded, since they would not explain differences in conjugation frequencies between isolates. Finally, only complete RM systems that were not present in the donor strains were retained for each recipient-donor combination, since the donor would be conferring protection to pOXA-48 against equivalent systems of the recipient. Complete type I systems comprise a restriction enzyme (REase), a methylase (MTase) and a specificity (S) subunit. Type II systems include at least a REase and a MTase. No type III systems were detected, and type IV systems are normally composed of one or two proteins. Donor and recipient systems were regarded as similar if they shared the same recognition sequence or if a BLASTp alignment of the enzymes followed the previous criteria of identity and alignment percentage per type of system. The same method was applied for comparing the final systems between recipient strains, and similar systems within each type were given the same letter code identifying the RM system subtype (Fig. S15, Table S3).

Clustered regularly interspaced short palindromic repeats (CRISPRs) were identified in the nucleotide sequences of the recipient strains with the DefenseFinder webserver. The spacer sequences were aligned to pOXA-48_K8 (MT441554) using the blastn-short function of BLASTn v2.9.0 (60) and hits were then filtered by percentage of alignment. Other defense systems were also detected with the DefenseFinder webserver and PADLOC v1.1.0 (database v1.3.0, (64)).

Plasmid incompatibility was investigated by identifying plasmid replicons using ABRicate v1.0.1 with the PlasmidFinder database.

Secretion systems were searched using the TXSScan models of MacSyFinder v1.0.5 (62, 65), selecting the option “ordered_replicon” and linear topology. Only complete systems were considered.

Capsular systems were identified using two software. First, *Klebsiella* capsules, which belong to the Wzx/Wzy-dependent group I capsular systems, were identified with Kaptive v2.0.0 (66), using the K locus primary reference and default parameters. The presence of *Klebsiella* capsules was assigned when the confidence level of the matches was above “good”, as recommended by the authors. Other capsular systems from groups I (Wzx/Wzy-dependent), II and III (ABC-dependent), IV, synthase-dependent (cps3-like and hyaluronic acid) and PGA (Poly-Y-d-glutamate) were searched using MacSyFinder v1.0.5 with the CapsuleFinder for diderms models, indicating “ordered_replicon” as database type and linear topology (62, 67). Only systems reported as complete were considered. In the cases where *Klebsiella* capsular loci were also identified by CapsuleFinder as group I capsules, analyses were performed with the Kaptive output. To assess the similarity of the different sets of capsular loci between strains, a weighted gene repertoire relatedness (wGRR) score was calculated as in (68). For this, significant sequence similarities (e-value <10^-4^, identity ≥35%, coverage ≥50%) among all pairs of proteins in the capsular loci were searched using MMSeqs2 (Nature Biotechnology release, August 2017, (69)) with the sensitivity parameter set to 7.5. The best bi-directional hits between pairs of capsule loci sets were then used to compute the wGRR as:

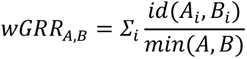

where *id(A_i_, B_i_)* is the sequence identity between each pair *i* of homologous proteins in *A* and *B*, and *min(A, B)* is the number of proteins of the capsule loci set encoding the fewest proteins (*A* or *B*). The wGRR takes values between 0 (no homologous proteins between capsule loci sets) and 1 (all genes of the smallest capsule loci set have an identical homologous in the loci set of the other strain), representing the fraction of homologs in the smallest of the two capsule loci set weighted by their sequence similarity. Protein sequences of the *Klebsiella* capsular loci were obtained from Prokka annotations of the operon nucleotide sequences output by Kaptive. All proteins between the borders of CapsuleFinder complete systems were included.

### Analysis of capsular systems in a database

The database comprised the genomes of 730 *K. pneumoniae* and 1,585 *E. coli* downloaded from the NCBI RefSeq database of high quality complete non-redundant genomes (retrieved from ftp://ftp.ncbi.nlm.nih.gov/genomes/refseq/ on March 2021; Table S4) (70). pOXA-48-carrying strains were identified in the RefSeq database when the best, largest BLASTn hit between the sequence of pOXA-48_K8 (MT441554) and the target plasmid had >95% identity and >10 Kb alignment, and when the plasmid sequence contained the pOXA-48 IncL replicon and the *bla*_OXA-48_ gene as detected by ABRicate with the PlasmidFinder and ResFinder databases. The plasmids of 6 *E. coli* and 46 *K. pneumoniae* strains met this criteria, which had lengths between 50.6-74.2 Kb (average 64.0 Kb; Table S4). Kaptive and CapsuleFinder were used as previously described to search for *Klebsiella*-derived capsules and capsular systems, respectively, and to discard non-capsulated strains (15 *K. pneumoniae* strains).

Associations between the presence of *Klebsiella*-derived capsules and pOXA-48- or plasmid-carriage were analyzed building 2×2 contingency tables. Significant interactions (*P* < 0.05, Fisher exact test) were then corrected for phylogenetic dependency, i.e. the tendency of closely related strains to share the same traits. For this, a set of 128 HMM profiles (Pfam release 35, Table S5) of conserved bacterial single copy genes, described in (71) and curated in (72), were identified in the RefSeq strains and an *Enterobacter cloacae* outgroup (accession number GCF_003204095.1) using HMMER v3.3 (option--cut_ga) (73). Hits were filtered by score using the cutoffs reported in (65), resulting in 127 gene markers present in >90% of strains. Protein sequences of each family were aligned with MAFFT v7.453 (option--auto) (74) and alignments were trimmed with trimAl v1.4.rev15 (75). IQ-TREE v1.6.12 (76) was used to infer two phylogenetic trees from the concatenated alignments (the first including all capsulated strains, and the second including only *E. coli* strains, both with the *E. cloacae* outgroup) with best evolutionary model selection and 1000 ultrafast bootstrap. Trees were rooted and rescaled to a total height of 1. Phylogenetic logistic regression for each significant association was performed with the function *phyloglm* from the *phylolm* v2.6.4 R package (77), fitting a logistic MPLE model with 100 independent bootstrap replicates.

### Mathematical model and simulations of polymicrobial communities

A detailed description of the population dynamics model and the computational simulations can be found in the Supporting Information (SI Text). All code and data used is available at https://github.com/ccg-esb/EvK.

### Statistical analyses

All statistical analyses were performed with R v4.2.2 (www.R-project.org). Packages *ggplot2* v3.3.6, *ggpubr* v0.4.0 *pheatmap* v1.0.12, *RColorBrewer* v1.1-3, *ggsignif* v0.6.3 and *tidyverse* v1.3.1 were used for data manipulation and representation. R base packages were used for statistical tests. Normality of the data was assessed with the Shapiro-Wilk test. To analyze the differences in initial (PF) and final (TC) MIC of antibiotics and PCN between *E. coli* and *Klebsiella* spp. a Wilcoxon sum-rank test was used. To analyze the differences between the members of each recipient couple in the pairwise conjugation experiments (Shapiro-Wilk normality test, *P* < 0.01 in all cases), the non-parametric Wilcoxon paired signed-rank test was used. To determine differences between receptor species in the classical and pooled recipient conjugation experiments, as well as in *in vivo* conjugation assays (Shapiro-Wilk normality test, *P* < 0.01 in all cases), the non-parametric Kruskal-Wallis test was performed. Correlations were performed with the Spearman’s rank test. Phylogenetic analyses were performed with *ape* v5.6-2, *phytools* v1.0-3 and *phylolm* v2.6.4 packages.

### Data and code availability

Sequencing data generated in this study is available in the Sequence Read Archive (SRA) under the BioProject <u>PRJNA803387</u>. The R-GNOSIS sequences were generated in (32) and a subset of strains were selected as in (36) (see Methods). All source data is provided as Supporting Information. The code generated during the study along with detailed bioinformatic methods is available at https://github.com/LaboraTORIbio/super-sinks and https://github.com/ccg-esb/EvK.

## Acknowledgements

This work was supported by the European Research Council (ERC) under the European Union’s Horizon 2020 research and innovation programme (ERC grant no. 757440-PLASREVOLUTION) and by the Instituto de Salud Carlos III (PI19/00749) cofunded by the European Development Regional Fund ‘A way to achieve Europe’. The R-GNOSIS project received financial support from the European Commission (grant no. R-GNOSIS-F P7-HEALTH-F3-2011-282512). C.U. was supported by a grant from MICINN (PID2020-120292RB-I00). We appreciate Andrea Fernandez Duque’s contribution in creating the visual illustrations.

## Notes

https://github.com/LaboraTORIbio/super-sinks/

https://github.com/ccg-esb/EvK/

### Competing Interest Statement

The authors have declared no competing interest.

### Summary of Updates

We have substantially modified the manuscript to underline that we study antimicrobial resistance level and conjugation permissiveness associated with plasmid pOXA-48. We now avoid the generalization towards vertical/horizontal transmission dynamics. We have significantly improved our mathematical model, and we provide now a step-by-step description of this model in several pages of the Supporting Information (pages 2-12). We believe that this model provides key advantages compared with those form previous studies. Namely, the introduction of three different variables per strain with experimentally determined values: fitness in the absence of antibiotics (with and without pOXA-48), conjugation permissiveness, and resistance levels. Supplemental files are also updated.

