## Supporting Information for "Antimicrobial resistance level and conjugation permissiveness shape plasmid distribution in clinical enterobacteria"

### Supporting Information Text

### 1. Plasmid population dynamics model

This supplementary text outlines a mathematical model designed to study the plasmid dynamics within bacterial populations under the influence of antibiotic treatment. This exploration is conducted via computational simulations, designed to understand the behavior of plasmid-bearing populations in response to antibiotics under varied environmental conditions. Our model incorporates key elements such as resource-limited growth, antibiotic-induced killing, plasmid transfer and segregational loss. Through simulation of different population structures and environmental regimes, we study how these factors modulate the distribution of plasmids within the community and identify successful plasmid-host associations.

**Modeling bacterial growth.** We consider that growth of each bacterial strain is determined by several key parameters: the maximum uptake rate ($V$), the half-saturation constant ($K_m$), and a resource conversion coefficient ($\rho$). These parameters define the Monod term, which encapsulates the fundamental growth kinetics of bacterial strains. The uptake function $U(S)$, represented by the equation $U(S) = \frac{V \cdot S}{S+K_m}$, determines how the population consumes the limiting resource ($S$). The half-saturation constant $K_m$ represents the resource concentration at which the uptake rate is half of its maximum, hence providing a measure of the affinity of the bacteria for the resource. Previous studies have established that it is challenging to simultaneously parametrize the parameters $V$ and $K_m$ from experimental data, so we will use $V/K_m$, a structurally identifiable ratio commonly known as the 'specific affinity'(1). The other parameter relevant for the growth dynamics is the resource conversion coefficient $\rho$, which represents the efficiency with which cells convert the consumed resources into new biomass, determining the bacterial population's growth rate per unit of resource consumed. The growth function can thus be expressed as $G(S;\psi) = \rho \cdot U(S;\psi)$, with $\psi = (\rho, V/K_m)$ representing a vector of parameters that characterize the growth dynamics of a particular bacterial strain.

**Modeling the dynamics of plasmid-bearing and plasmid-free populations.** Plasmids replicate independently of the bacterial chromosome and therefore a bacterial population may be divided into two distinct subpopulations: plasmid-bearing cells ($B_p$) and plasmid-free cells ($B_\emptyset$). The presence or absence of plasmids within these subpopulations can significantly impact their overall behavior and response to environmental stressors, such as antibiotics. In our model, we assign distinct growth kinetic parameters to $B_p$ and $B_\emptyset$. Adjusting these parameters enables our model to capture a spectrum of fitness effects associated with plasmid-bearing, ranging from costs to benefits(2). It's important to note that although fitness costs can potentially be compensated over time(3), we treat them as constants in our model. This simplification is justified as our study is primarily focused on the ecological dynamics, replicating short-term experiments. Detailed investigation of evolutionary dynamics of the plasmid-host association itself is beyond the scope of this paper and could be an avenue for future research.

**Modeling plasmid segregation.** In addition to growth dynamics, our model accounts for the intrinsic processes occurring within plasmid-bearing bacterial cells. One crucial aspect is the segregational loss of plasmids, a random event that occurs during bacterial cell division. As a plasmid-bearing cell divides, it distributes its plasmids to its two offspring cells. Given the probabilistic nature of this distribution, there are instances when one of the offspring cells may not inherit a plasmid, thereby converting it into a plasmid-free cell. This stochastic event, denoted as 'segregational loss', represents an essential dynamic contributing to the overall plasmid count within the bacterial population and can lead to the reduction of plasmid-bearing cells in the absence of positive selection for plasmid-encoded genes. The rate at which this loss occurs is captured by a parameter $\lambda$.

The rate of segregational loss can depend on various factors. If plasmids were to segregate randomly upon division, we could model the segregational loss rate as a function of the plasmid copy number (PCN) in the cell(4). However, the plasmid we study in our model, pOXA-48, features an active partitioning system. This system aids a more balanced plasmid distribution among daughter cells during division, thus reducing the segregation loss rate. Plasmids with such a system typically have segregation loss rates between 0.1% and 0.2%(5, 6). In particular, we assume a constant segregation rate $\lambda = 0.002$ for all strains.

**Modeling plasmid conjugation.** The model incorporates the horizontal transfer of plasmids between cells via conjugation. We model the net change in the plasmid-bearing bacterial population due to conjugation events, taking into account the densities of both the plasmid-free ($B_\emptyset$) and plasmid-bearing ($B_p$) populations. In our well-mixed environment, we assume the principle of mass action, which in this context means that the probability of two cells encountering each other and engaging in conjugation is proportional to the densities of both the plasmid-free and plasmid-bearing subpopulations. We incorporate the rate of plasmid transfer, measured in conjugation events per donor cell per hour, by introducing a parameter that quantifies the propensity of a bacterial cell to accept a plasmid from another cell. This introduces a parameter, 'permissiveness' that we denote with $\gamma$, to measure a bacterial cell's propensity to accept a plasmid from another cell.

**Modeling susceptibility to antimicrobial drugs.** Our model incorporates the impact of antibiotic resistance on bacterial populations. Specifically, it represents the antibiotic concentration at any given time $t$ as $A(t)$. We are dealing with bactericidal antibiotics, such as beta-lactam antibiotics, that are subject to enzymatic degradation. The model includes an inactivation rate for antibiotic molecules, which is assumed to be proportional to the total bacterial population and the environmental antibiotic concentration. We consider the rate of degradation to be higher for plasmid-bearing cells, $\alpha_p > \alpha_\emptyset \geq 0$. The antibiotic-induced death rate in the model is governed by the antibiotic concentration at any given time, the size of the bacterial subpopulation, and a kill rate given by $\kappa = 1/(\hat{\kappa} * \text{MIC}_{max})$, where $\hat{\kappa}$ denotes the strain's resistance coefficient and $\text{MIC}_{max}$ a normalizing constant denoting the maximum drug used. In this context, $\hat{\kappa}$ is a measure of the bacterial strain's inherent capacity to resist the antibiotic - a larger $\hat{\kappa}$ implies greater resistance.

**Plasmid population dynamics model.** Altogether, our model captures the dynamics of a bacterial population composed of two distinct subpopulations: plasmid-bearing ($B_p$) and plasmid-free ($B_\emptyset$) cells. Both subpopulations compete for a limiting resource, $S$, and are susceptible to an antibiotic, $A$. The interplay between vertical and horizontal plasmid transmission routes via segregational loss and conjugation, respectively, plays a significant role in shaping the plasmid distribution and persistence within these subpopulations over time.

We describe the system through a set of differential equations, which encapsulate the temporal evolution of the system state $\bar{x} = (S, A, B_\emptyset, B_p)$, and that can be written as:

$$\frac{dS}{dt} = -\left(U(S;\psi_\emptyset)\cdot B_\emptyset + U(S;\psi_p)\cdot B_p\right), \quad [1]$$

$$\frac{dA}{dt} = -A\cdot(\alpha_\emptyset B_\emptyset + \alpha_p B_p), \quad [2]$$

$$\frac{dB_\emptyset}{dt} = (G(S;\psi_\emptyset) - \kappa_\emptyset A)\cdot B_\emptyset + \lambda\, G(S;\psi_p)\cdot B_p - \gamma\, B_\emptyset\, B_p, \quad [3]$$

$$\frac{dB_p}{dt} = (1 - \kappa_p A - \lambda)\, G(S;\psi_p)\cdot B_p + \gamma\, B_\emptyset\, B_p, \quad [4]$$

The initial conditions are determined by the starting resource concentration, which is set to $S_0 = 1$ throughout our study, and the environmental drug concentration, $A_0 \geq 0$. The initial densities of $B_p(0)$ and $B_\emptyset(0)$ are given by the specific experimental setup being simulated.

**Numerical examples of plasmid population dynamics.** To illustrate the plasmid population dynamics, we will simulate a drug-free scenario by setting the initial environmental antibiotic concentration as $A(0) = A_0 = 0$. Therefore the simulation's initial conditions are defined by the variable $y_0 = (S_0, A_0, 0, B_0)$, which represents the initial substrate ($S_0$) and drug concentrations ($A_0$), and the initial bacterial density, $B_0$. Fig. S1A shows an example where all cells in the population are plasmid-bearing with an initial density of $B_0 = 1 \times 10^6$. Note that the population contains a stable subpopulation of plasmid-free cells produced by segregational loss. In comparison, Fig. S1B depicts a population of cells without plasmids, growing until their available resources are depleted.

Fig. S1C shows an example whereby the environmental antibiotic concentration is set to $A = 1$, indicating that the bacterial populations are now subject to antibiotic pressure. Under these conditions, the plasmid-bearing population outperforms the plasmid-free cells due to their enhanced resistance to the antibiotic. The survival advantage of the plasmid-bearing cells is depicted in the resulting plot, showing the role of plasmids by allowing the $B_p$ cells to proliferate when $A > 0$. Also note that a small fraction of plasmid-free cells is maintained through segregational loss. In comparison, Fig. S1D depicts a similar numerical experiment involving only plasmid-free cells, indicating that the antibiotic results in the complete eradication of the entire $B_\emptyset$ population.

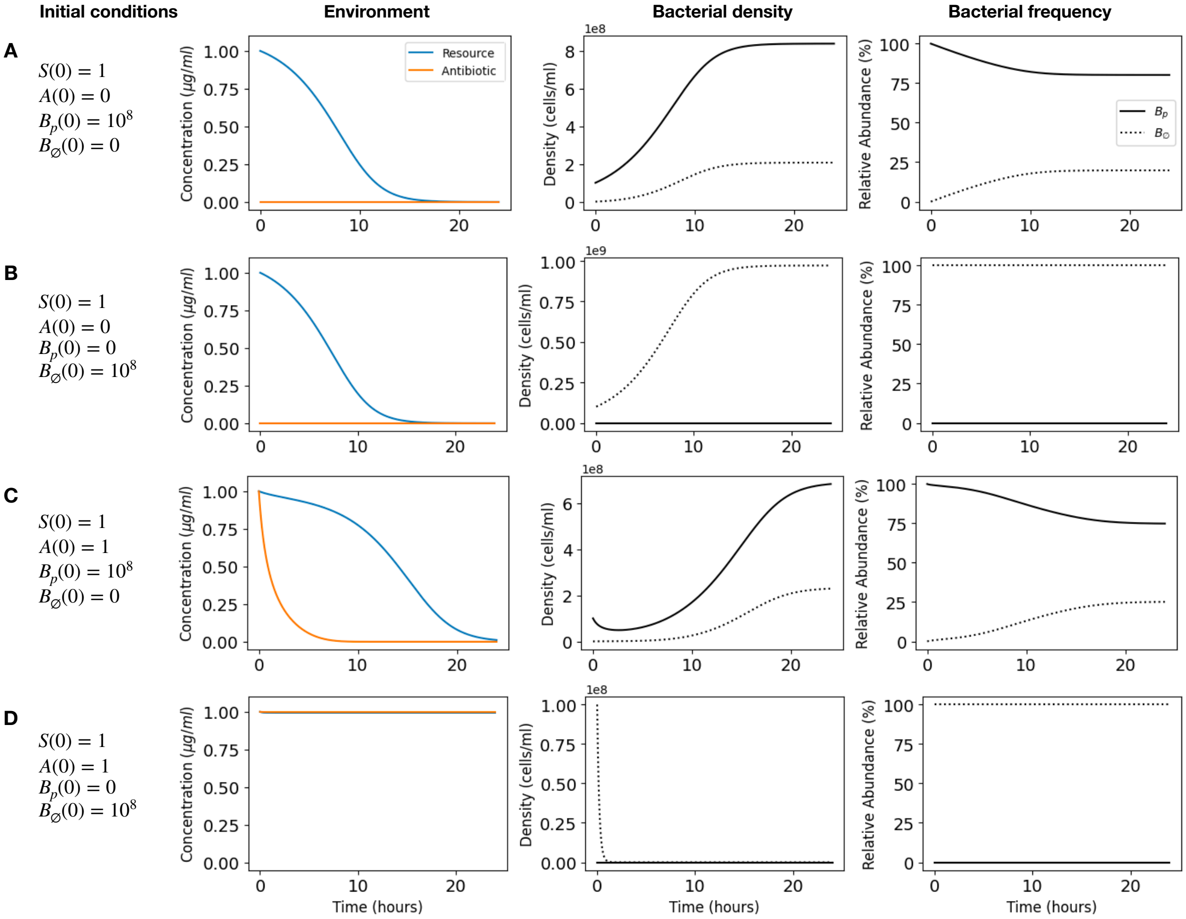


**Fig. S1. Numerical examples of plasmid population dynamics.** (A) This panel presents the dynamics of a drug-free environment ($A(0) = 0$), initially populated with plasmid-bearing bacteria ($B_p(0) = 1 \times 10^8$). The columns from left to right respectively show: resource and antibiotic concentrations (blue and orange lines, respectively), bacterial density illustrating the dominance of plasmid-bearing population and the emergence of plasmid-free segregants, and frequency of bacterial populations, which highlights the plasmid-bearing majority and low-frequency segregants. (B) The panel illustrates dynamics in a drug-free environment ($A(0) = 0$) initially inoculated with plasmid-free bacteria ($B_\emptyset(0) = 1 \times 10^8$) only. The columns from left to right display: resource and antibiotic concentrations, bacterial density showing the growth of the plasmid-free population, and frequency, which reveals a complete predominance of plasmid-free bacteria due to the absence of any initial plasmid-bearing population. (C) Population dynamics in an environment supplemented with antibiotics ($A(0) = 1$) and initially inoculated with plasmid-bearing bacteria ($B_p(0) = 1 \times 10^8$). Note the degradation of the antibiotic and the prevalence of the plasmid-bearing bacteria and the presence of plasmid-free cells due to segregation. (D) The panel portrays dynamics in an antibiotic environment ($A(0) = 1$), initially populated with plasmid-free bacteria ($B_\emptyset(0) = 1 \times 10^8$), showing the eradication of the plasmid-free population.

### 2. Model parametrization

To parameterize our model, we extract data from experimental results obtained from growing each bacterial strain in isolation. These parameters encompass aspects such as growth kinetics, susceptibility to antibiotics, and conjugative permissiveness. The goal of this parameterization process is to determine a parameter set capable of simulating the ecological dynamics of mixed co-cultures. This enables the execution of *in silico* competition experiments under designated environmental conditions.

**Growth-kinetic parameters.** In a prior study(2), we employed the Metropolis-Hastings Markov chain Monte Carlo (MCMC) approach to extract maximum likelihood estimates for growth parameters within our ecological model. Using this method, we were able to determine the specific affinity ($V/K_m$) and the resource conversion rate ($\rho$) for every bacterial strain from isolated growth curves, encompassing both plasmid-bearing and plasmid-free cells. Through model fitting to individual growth curves, we generated a parameter vector $\psi = (\rho, V/K_m)$ for each strain (Fig. S2A-B). Fig. S2C-D shows the function $U(S)$ evaluated with these parameters estimated for each strain. This demonstrates a wide distribution of fitness effects, which is expressed in terms of overlapping uptake rates between plasmid free and plasmid bearing cells (2).

To evaluate the robustness of our model, we considered various prior distributions (uniform, lognormal, beta, and gamma). The results from these different distributions consistently produced similar maximum likelihood estimates. Moreover, by employing a data cloning algorithm (7, 8), we were able to validate the identifiability of our parameters. This was reflected in the convergence of the marginal posterior distribution to the maximum likelihood estimate with the increase in the number of clones.

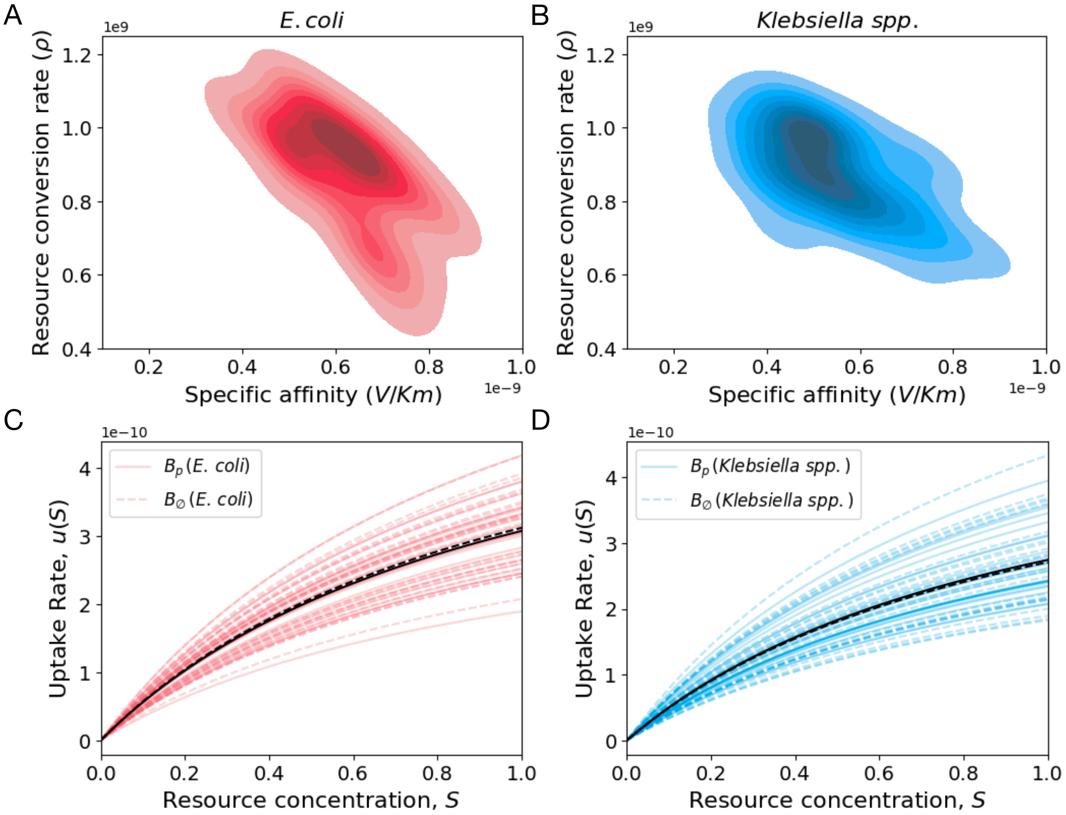


**Fig. S2. Parametrization of growth-kinetic parameters.** (A) Kernel Density Estimation (KDE) plot for *E. coli* strains, visualizing the joint distribution of specific affinity ($V/K_m$) and resource conversion rate ($\rho$). The filled contour plot was generated using a Gaussian Kernel Density Estimation. The color map represents the density of data points, with darker shades of red indicating a higher density of observations. (B) KDE plot for *Klebsiella spp.* strains, constructed in a manner similar to (A). The variations in shades of blue represent the relative density of observations. (C) Plot of the resource uptake rate, $U(S)$, as a function of resource concentration, $S$, for *E. coli* strains. Each red line represents a different *E. coli* strain, with plasmid-bearing and plasmid-free strains distinguished by line type (solid for $B_p$ and dashed for $B_\emptyset$). The solid black line represents the mean uptake rate for plasmid-bearing *E. coli* strains, and the dashed black line represents the mean for plasmid-free *E. coli* strains. (D) Each line illustrates the uptake rate as a function of resource concentration for different *Klebsiella spp.* strains in our collection. The solid and dashed black lines represent the mean uptake rates for plasmid-bearing and plasmid-free *Klebsiella spp.* strains, respectively.

**Parametrizing susceptibility to bactericidal antibiotics.** Drug susceptibility of each strain plays a crucial role in determining how antibiotic concentration impacts the growth and survival of mixed bacterial populations. It also influences the dynamics of plasmid transfer as it can determine the selective advantage conferred by the plasmid in environments with varying antibiotic concentrations. In our model, we denote with $\kappa$ a parameter that measures a strain's resistance to the antibiotic, with higher values indicating increased resistance.

Experimentally, we exposed the population to doubling antibiotic concentrations of amoxicillin/clavulanic acid to identify the lowest concentration of the antibiotic at which bacterial growth is effectively inhibited, a quantity referred to as the Minimum Inhibitory Concentration (MIC) and expressed in units of $mg/L$. Fig. S3A illustrates the Parameter $\kappa$ can be inferred by fitting *in silico* by producing dose-response experiments and comparing the MIC obtained with the simulations with the MIC values obtained experimentally. We then performed a grid algorithm to identify $\kappa^*$, the parameter that best approximates the experimentally-determined MIC. If we repeat this numerical experiment for the plasmid-bearing population, we find that the population can still exhibit growth at high drug concentrations.

Fig. S3B Fig. S3C show theoretical dose-response curves obtained by plotting, respectively, the final bacterial density for plasmid-free and plasmid-bearing populations as a function of the drug concentration.

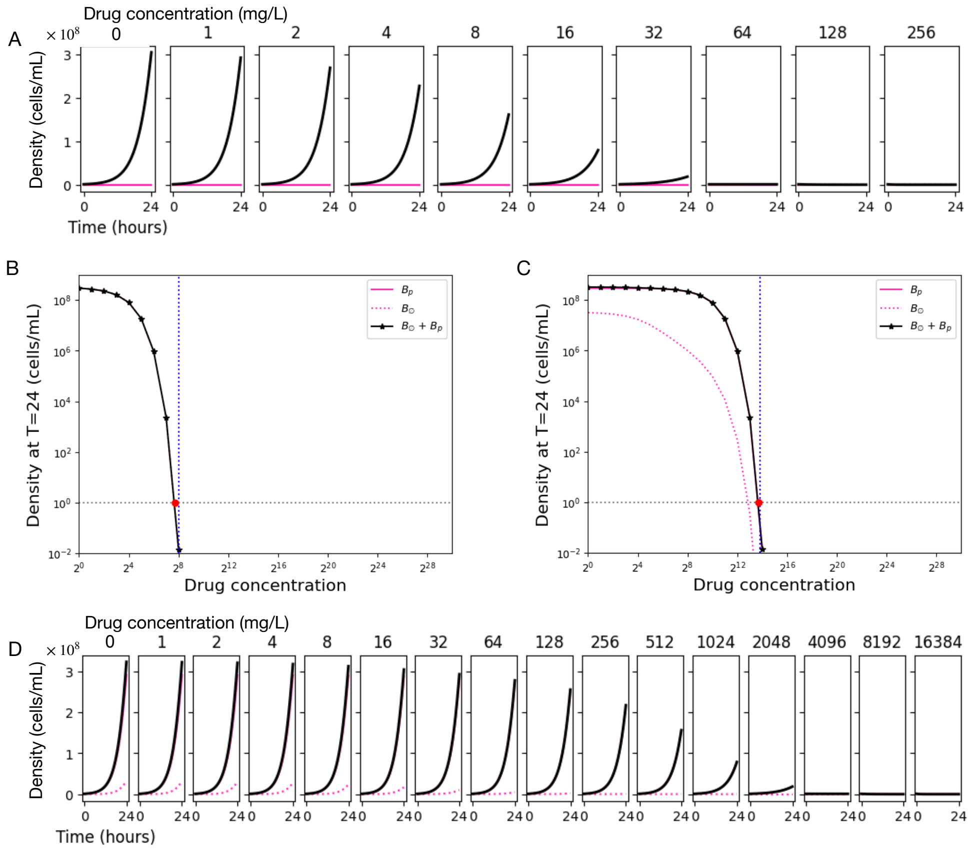


**Fig. S3. Parametrizing susceptibility to antibiotics** (A) This panel presents a theoretical dose-response curve for a single bacterial strain under a range of antibiotic concentrations. Each plot represents a simulation run at a specific antibiotic concentration (in units of mg/L), increasing in a doubling series from left to right. The total bacterial density over time is depicted by the black line. As expected, bacterial growth is suppressed in a dose-dependent manner by the antibiotic, with higher concentrations leading to greater suppression. (B) The final bacterial density at $T = 24$ is plotted as a function of drug concentration. The blue dotted line represents the experimentally determined MIC for the strain. The theoretical MIC, shown as the red dot, is obtained by fitting an *in silico* model to the data by identifying the optimal parameter, $\kappa^*$, that best describes the experimental dose-response experiment. (C) and (D) are similar to (B) and (A), but represent a strain bearing a plasmid conferring antibiotic resistance. Note the shift in the dose-response curve compared to the plasmid-free strain, reflecting the increased survival and growth of the plasmid-bearing strain at higher antibiotic concentrations.

### 3. Plasmid Dynamics in Multistrain Populations

**Multistrain population dynamics model.** We extended the plasmid population dynamics model to include a conjugative plasmid spreading in a population composed of multiple bacterial strains. These cells compete for a single, limiting resource within a homogeneous environment. Our model assumes strictly competitive interactions between strains, without any metabolic exchanges. Moreover, our model enables the simulation of horizontal plasmid transfers between different strains through conjugation. By incorporating multistrain dynamics, we create a framework to explore the population dynamics of plasmids within diverse bacterial communities of $M$ different strains, both plasmid-free and plasmid-bearing,under a variety of selective pressures.

In the multistrain model, we consider that each bacterial strain $i$ is represented by the densities of the plasmid-bearing and plasmid-free subpopulations, denoted as $B_{pi}$ and $B_{\emptyset i}$, respectively. Thus, we define the bacterial community at a given time $t$ as a vector: $\mathbf{B}(t) = (B_{p1}(t), B_{p2}(t), ..., B_{pM}(t), B_{\emptyset 1}(t), B_{\emptyset 2}(t), ..., B_{\emptyset M}(t))$.

Each cell consumes a limiting resource at rates determined by its unique growth kinetic parameters.

$$\frac{dS}{dt} = -\sum_{i=1}^{M}(U_{pi}(S)B_{pi} + U_{\emptyset i}(S)B_{\emptyset i}).$$

In our model, we assume that antibiotic molecules degrade at a rate proportional to the total bacterial population and the environmental drug concentration,

$$\frac{dA}{dt} = -A\sum_{i=1}^{M}(\alpha_{pi}B_{pi} + \alpha_{\emptyset i}B_{\emptyset i}).$$

The dynamics of the plasmid-bearing subpopulation of strain $i$ depends on segregation and conjugation of the plasmid, as well as growth and death rates of both $B_{pi}$ and $B_{\emptyset i}$,

$$\frac{dB_{pi}}{dt} = G(S;\psi_{pi})\,(1-\kappa_{pi}A-\lambda)\,B_{pi} + \gamma_i\,B_{\emptyset i}\sum_{j=1}^{M}B_{pj}$$

.

Likewise, the plasmid-free subpopulation of strain $i$ follows the dynamics:

$$\frac{dB_{\emptyset i}}{dt} = G(S;\psi_{\emptyset i})\,(1-\kappa_{\emptyset i}A)\,B_{\emptyset i} + \lambda\,G(S;\psi_{pi})B_{pi} - \gamma_i\,B_{\emptyset i}\sum_{j=1}^{M}B_{pj}.$$

**Numerical example of a serial dilution experiment.** Serial dilution experiments are important tools in microbiology, helping to study how bacteria grow, survive, and behave under different conditions (9). These experiments follow a cycle: bacteria are grown in a culture, then diluted, and then allowed to grow again in fresh media. Our computer simulations reproduce this process using the mathematical model to describe the dynamics of multiple bacterial strains competing for a single limited resource ($S_0$) and exposed to antibiotic at concentration ($A_0$).

To mimic a situation where a plasmid is introduced into the population, we assume that all bacteria are initially without a plasmid, except one strain that carries a plasmid. So, if there are $M$ strains in the population, the starting density of each plasmid-free subpopulation at time $t = 0$ is set to $B_\emptyset(0) = B_0/M$, where $B_0$ is the total initial bacterial density (in our experiments, $B_0 = 10^6 - 10^8$), and the starting density of the plasmid-bearing subpopulations is set to zero. We then make the plasmid-carrying bacteria of a random strain equal to 0.01% of $B_0$, indicating the start of the plasmid spreading.

Starting from the second day, we simulate a daily 'serial dilution' process. At the start of each day, we calculate the bacterial densities from the end of the previous day, and then dilute by a factor of $d = 0.1$. This simulates the dilution event and sets the starting conditions for the new day. At the start of each day, we also restore the resource concentration to $S_0$ and add antibiotic with a concentration $A_0$. This cycle of growth, dilution, and regrowth continues for a set number of days (in our experiments, $N = 10$). For each drug concentration, the simulation tracks the density of each subpopulation over time (Fig S4).

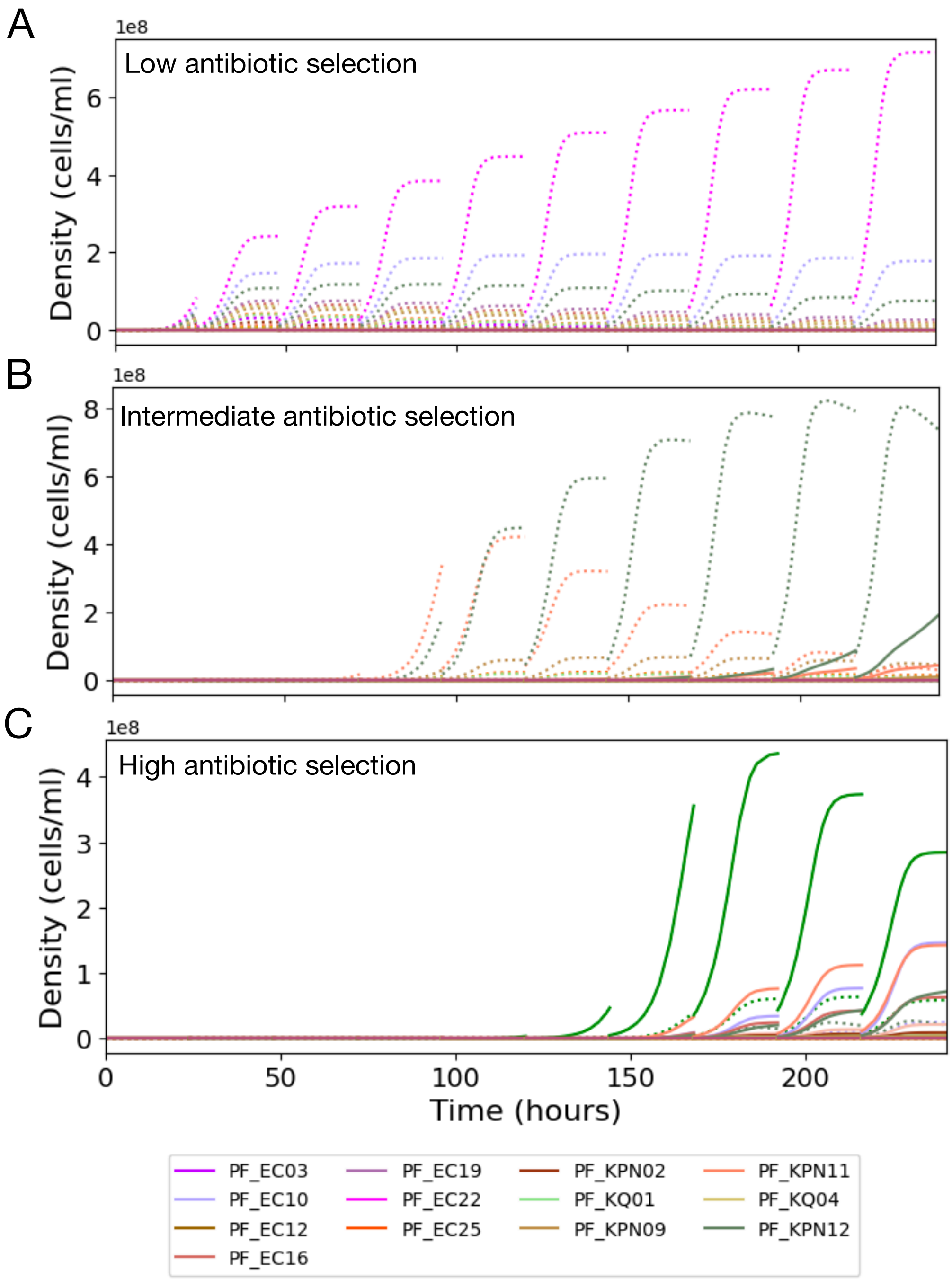


**Fig. S4. Numerical examples of serial transfer experiments** (A) The figure represents the density of each strain in the population as a function of time in a drug-free environment ($A = 0$ mg/L). Each unique color represents a different strain. Dotted lines represent plasmid-free bacteria, while solid lines denote plasmid-bearing bacteria. In the legend, only those strains that exhibited observable growth are displayed. (B) Density of each strain exposed to intermediate selective pressures ($A = 32$ mg/L). (C) At high antibiotic concentrations ($A = 1024$ mg/L), there is reduced growth and a strong selective pressure favoring plasmid-bearing bacteria.

**Computer experiment: Pairwise competition assays.** Our pairwise competition assays aim to investigate the competitive interactions between specific bacterial strains of *Klebsiella spp.* and *E. coli*. These simulations utilize parameters derived from experimental data. The simulations begin by initializing each strain with an equal population density, evenly split between plasmid-bearing and plasmid-free subpopulations. For every competing strain pair $i$, the initial density of plasmid-bearing cells at time $t = 0$ is set as $B_{pi}(0) = B_0/2$, and the plasmid-free density is $B_{\emptyset i}(0) = B_0/2$.

Subsequently, these populations are exposed to varying antibiotic concentrations, and the plasmid population dynamics are tracked over a 10-day experiment (Fig S5A). This approach enables us to estimate the proportion of stable plasmid occurrences within *Klebsiella spp.* and *E. coli* pairs across various drug concentrations. We observed that plasmid stability correlates with the strength of selection: *E. coli* predominantly hosts the plasmid under intermediate drug pressures, while plasmid-bearing *Klebsiella spp.* populations dominate under high drug levels (Fig. S5B).

By calculating the area under the curve (AUC) for each strain at every antibiotic concentration, we assess their combined growth and survival effects throughout the duration of the experiment. The strain with the higher AUC is considered to possess greater fitness under the specific antibiotic concentration. The results of all pairwise experiments between *E. coli* and *Klebsiella spp.* strains at different selective pressures are presented in Figs. S6 and S7.

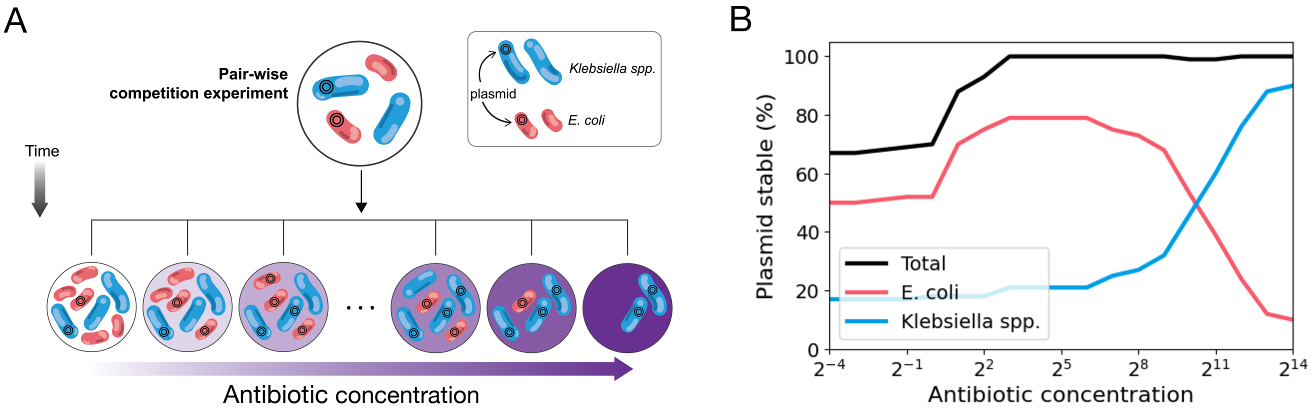


**Fig. S5. Computer experiment: Pair-wise competition experiment** A) Diagram representing a pairwise competition experiment conducted across a range of selective pressures. Starting with equal initial densities of both strains, each half comprised of plasmid-bearing and plasmid-free cells, our model allows us to forecast the distribution of strains and plasmids within the population under varying selective pressures. B) Proportion of stable plasmid occurrences within *Klebsiella spp.* and *E. coli* pairs across different drug concentrations. Plasmid stability correlates with selection strength: *E. coli* primarily hosts the plasmid at intermediate pressures, whereas plasmid-bearing *Klebsiella spp.* dominate at high drug levels.

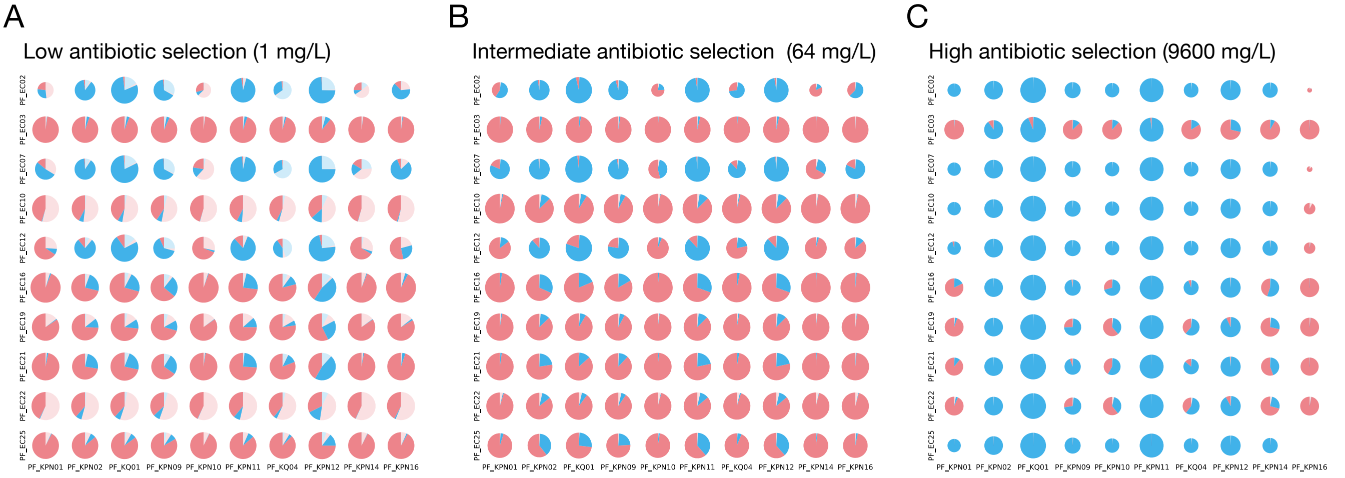


**Fig. S6. Species-level pairwise competition assays** Each circle represents a pairwise competition experiment between *E. coli* and *Klebsiella spp.* strains. Light colors denote plasmid-free bacteria, while dark colors indicate plasmid-bearing. Red is designated for *E. coli*, and Blue for *Klebsiella spp.* A) In conditions of low antibiotic selection (mg/L), there is a co-existence between plasmid-bearing and plasmid-free cells, with *E. coli* strains generally demonstrating a higher relative fitness. B) Under intermediate selection pressures, certain associations between *E. coli* and the plasmid prove particularly successful in maintaining the plasmid within the population. C) At high antibiotic selection levels, the population is primarily composed of plasmid-bearing *Klebsiella strains*, reflecting the high-pressure environment.

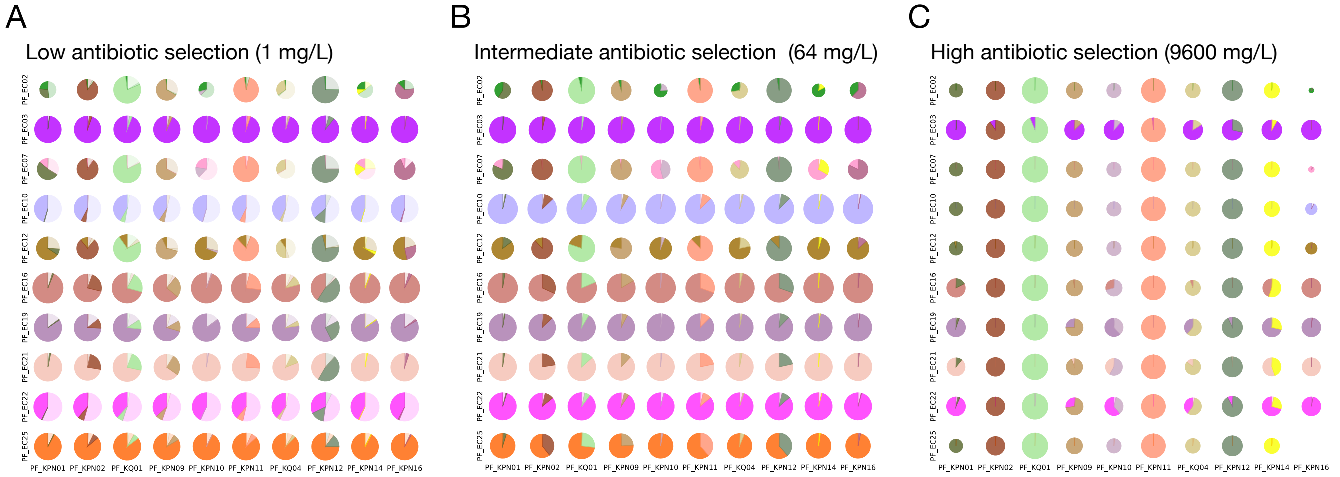


**Fig. S7. Strain-level pairwise competition assays** Each circle symbolizes a pairwise competition experiment between distinct *E. coli* and *Klebsiella spp.* strains. Now, colors are assigned uniquely to each strain, and both lighter and darker shades represent plasmid-free and plasmid-bearing states, respectively. A) At low antibiotic selection, coexistence between plasmid-free and plasmid-bearing cells is observed. Plasmid-bearing strains PF_EC25 and PF_EC03 (*E. coli*) display remarkable competitive fitness. B) Under intermediate selection pressure, plasmid-bearing *E. coli* strains dominate, indicating their advantage in maintaining the plasmid within these conditions. C) At high antibiotic selection, plasmid-bearing *Klebsiella spp.* strains, specifically PF_KQ01, PF_KPN11, and PF_KPN12, outcompete all *E. coli* strains, due to the lower resistance levels exhibited by the latter, leading to their dominance in the population. This underscores the heightened resilience of these *Klebsiella spp.* strains under intense antibiotic pressures.

**Computer experiment: Multistrain dose-response assay.** In our multistrain dose-response experiments, we study the impact of a range of amoxicillin/clavulanic acid concentrations on a co-culture of *Klebsiella spp.* and *E. coli* strains. These dose-response experiments comprise computational simulations initialized with a multi-strain bacterial community initially composed of 20 strains. We consider that only one member of the community carries the pOXA-48 plasmid, starting at a frequency of 0.01% of the total bacterial density Fig. S8A. Throughout the simulations, we monitor the density of each strain and the fraction of each strain carrying the resistance plasmid under different antibiotic concentrations. Fig. S8B illustrates the final density of each bacterial strain under varying levels of selective pressure, illustrating the antibiotic resistance fitness landscape for mixed bacterial populations.

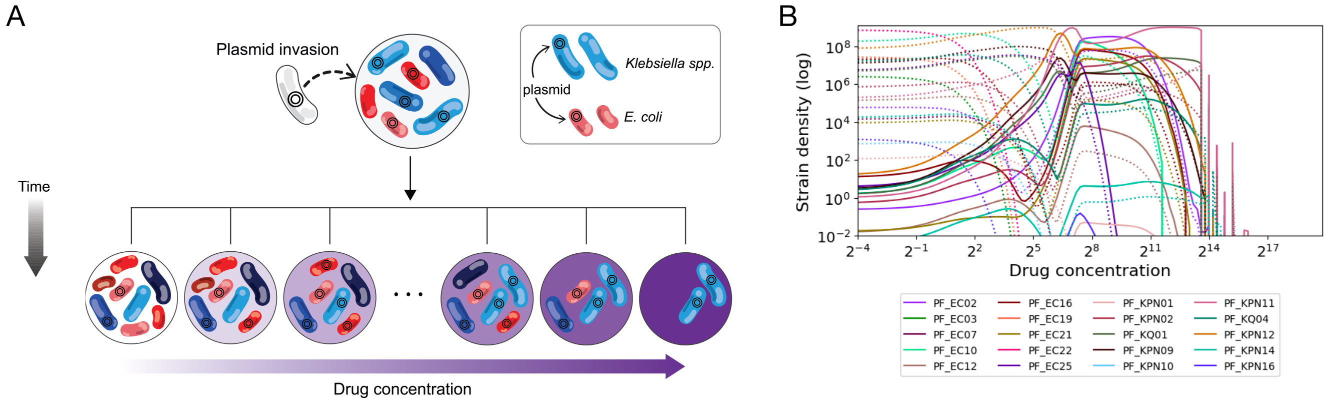


**Fig. S8. Computer experiment: Multistrain dose-response assay** A) Schematic representation of the multistrain dose-response numerical experiment: The initial community consists of 20 strains (10 *Klebsiella spp.* and 10 *E. coli*). These strains were used to parameterize the model based on data from cells grown in isolation. At t=0, the population is composed of equal densities of all plasmid-free strains and is invaded by a small fraction of cells carrying the plasmid belonging to a single strain. B) Through our model, we predicted the ecological dynamics and determined the final density (in log scale) of each strain after a 10 day experiment with a set daily drug concentration. Each strain's outcome is depicted by a unique color: dotted lines represent plasmid-free strains and solid lines indicate plasmid-bearing strains.

**Computer experiment: Iterative knock-out simulations.** To identify bacterial strains crucial for plasmid stability, we employed an iterative knock-out simulation approach. Sequentially, we removed strains from an initial community of 20 strains. The simulation was then re-initialized using the remaining strains, maintaining the original total bacterial density and the initial plasmid-bearing fraction, as in the multistrain dose-response assays (Figs. S9A). Fig. S9B shows the percentage of experiments where the final density above an arbitrary threshold for a range of drug concentrations and community sizes. Here we consider that a population survived if $\sum_1^M (B_{pi}(T) + B_{\emptyset i}(T)) > 1$.

Fig. S9C illustrate the number of times where removal of this strain resulted in the fraction of plasmid-bearing bacteria in the population below 50% after performing $1,000$ knockout experiments in environments with low antibiotic concentration ($A < 64$ mg/L). Interestingly, strains PF_EC03 (ST131), PF_EC16 (ST38), and PF_KPN11 (ST307) appear to be very relevant for maintaining plasmid stability in environments with low antibiotic concentration. In environments with high selective pressures ($A \geq 64$ mg/L), strains such as PF_KPN02 (ST485), PF_KQ01, and again PF_KPN11 (ST307) stand out as particularly critical for the dissemination and maintenance of plasmid pOXA-48 (Fig. S9D).

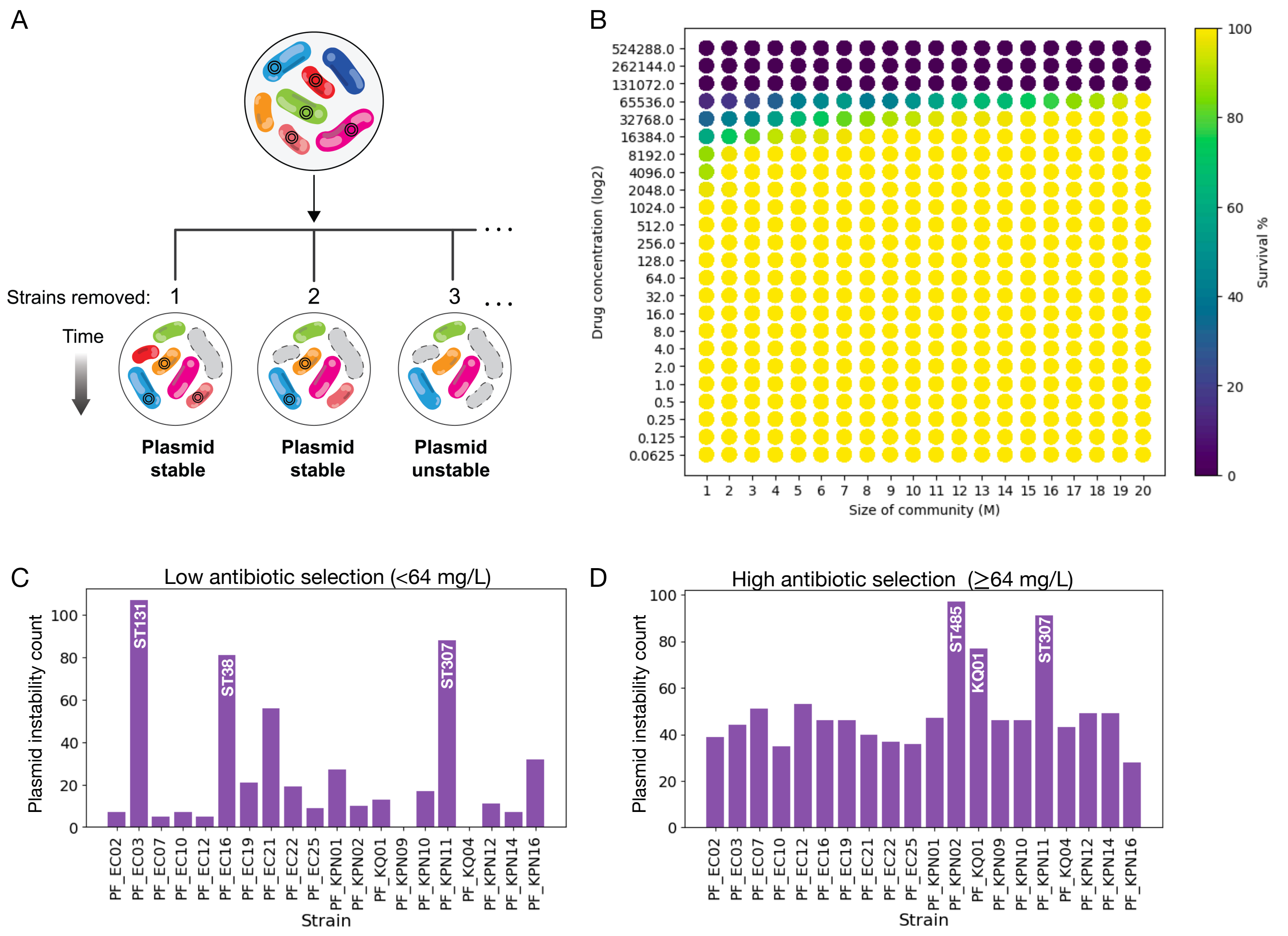


**Fig. S9. Strain knockout computational experiment** A) Schematic diagram illustrating the strain knockout computational experiment B) Fraction of experiments where the population density was above an arbitrary threshold for a range of drug concentrations and community sizes. Dark colors represent low probability of survival and light colors a high probability of survival. Note that increasing the size of the community results in an enhancement in the fraction of surviving populations. C) Bar chart illustrating the results of an iterative knock-out simulation in environments with low antibiotic selection ($A < 64$ mg/L). D) Bar chart showing the stabilizing effect of each strain under high antibiotic concentration settings ($A \geq 64$ mg/L), based on the iterative knock-out simulation.

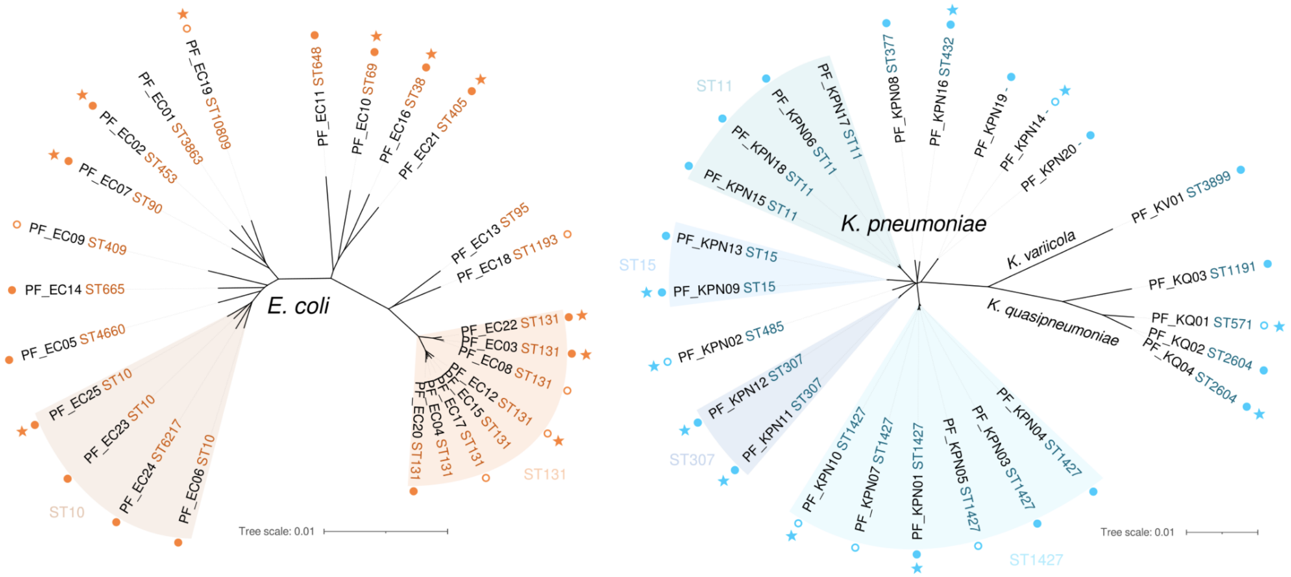


**Fig. S10.** Unrooted phylogeny of *E. coli* (left, $n = 25$) and *Klebsiella spp.* (right, $n = 25$) pOXA-48-free (PF) strains. Branch lengths represent mash distances between the whole-genome assemblies of the strains. Groups of strains belonging to the same multilocus sequence type (ST) are shaded (note that *E. coli* ST6217 is part of the ST10 group). Filled circles indicate strains for which transconjugants (TC) with an isogenic pOXA-48 could be obtained and were therefore used to determine minimum inhibitory concentrations (MIC) to antibiotics and pOXA-48 copy number (*E. coli* $n = 15$, *Klebsiella spp.* n=18). Empty circles indicate that TC could be obtained but harbored a mutated pOXA-48 plasmid. PF strains used as recipients in conjugation experiments are marked with a star.

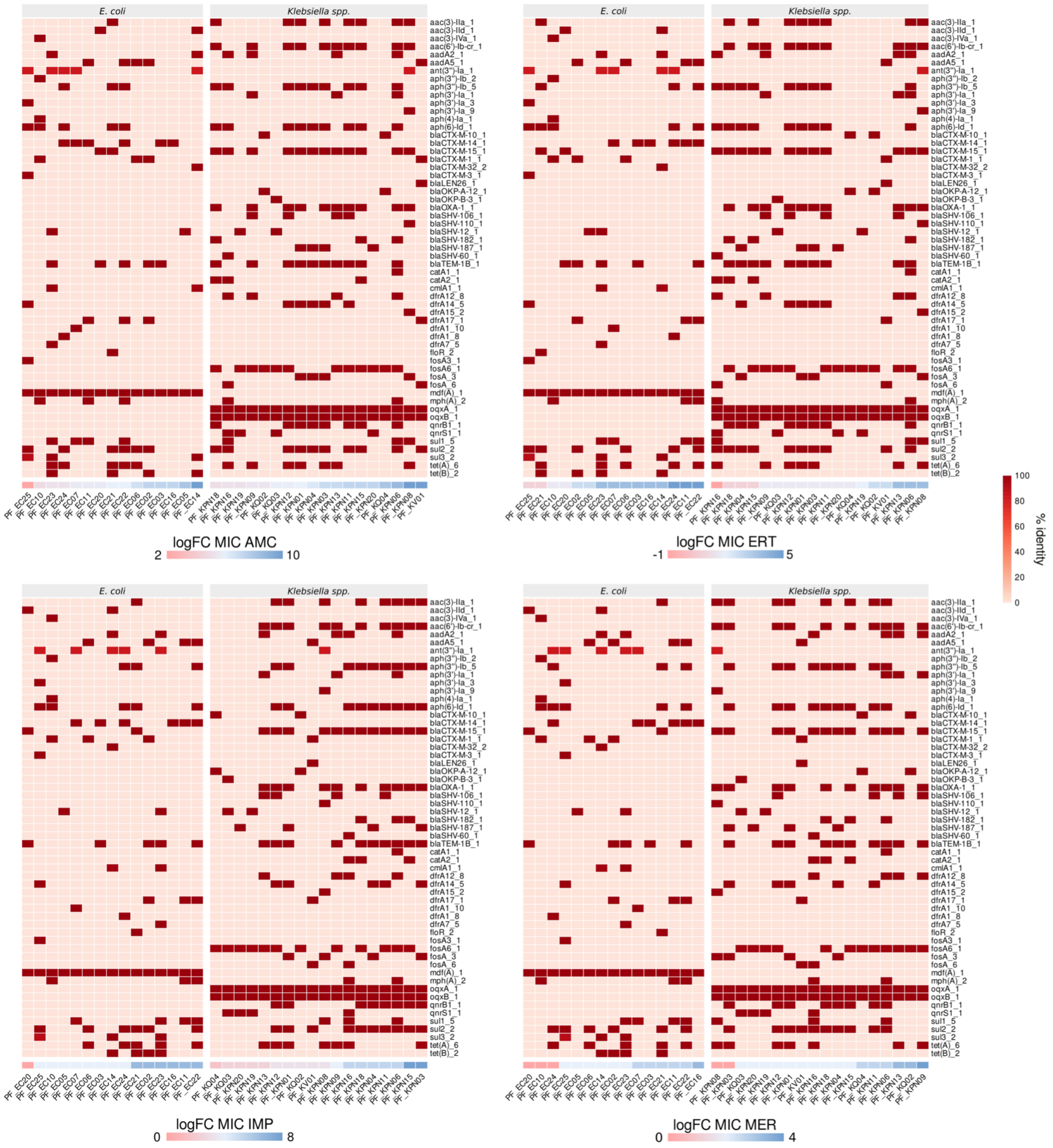


**Fig. S11.** Antimicrobial resistance (AMR) genes encoded by the pOXA-48-free (PF) strains used to determine minimum inhibitory concentrations (MICs) to antibiotics. The heatmap shows presence (colored by percentage of identity to the BLAST hit in the ResFinder database; see Methods) and absence (values of 0) of the AMR genes. In each of the four panels, the PF strains are ordered by fold change (FC) in MIC to the corresponding antibiotic when acquiring pOXA-48 (top left, amoxicillin/clavulanic acid, AMC; top right, ertapenem, ERT; bottom left, imipenem, IMP; bottom right, meropenem, MER).

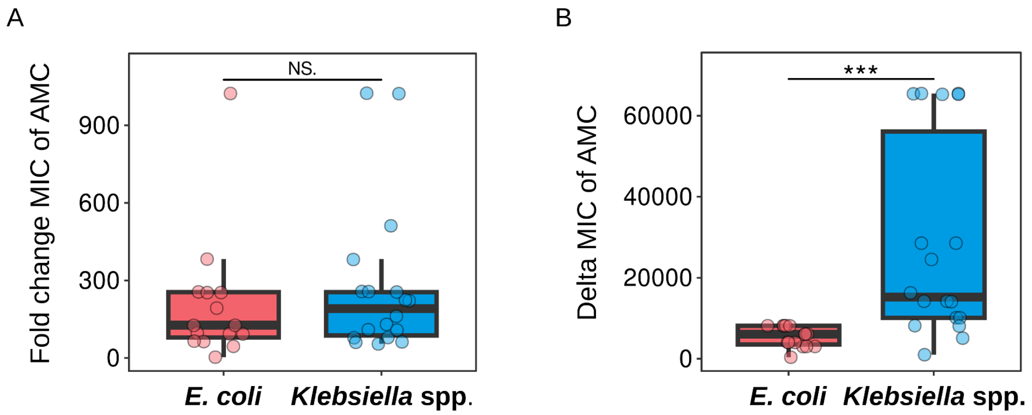


**Fig. S12.** Change in minimum inhibitory concentration (MIC) of amoxicillin/clavulanic acid (AMC) between pOXA-48-carrying and pOXA-48-free strains. (A) Fold change in MIC of AMC. (B) Difference (delta) in MIC of AMC between plasmid-carrying and plasmid-free pairs of strains. Horizontal lines inside boxes mark median values, the upper and lower hinges correspond to the 25th and 75th percentiles, and whiskers extend to 1.5 times the interquartile range. Dots indicate individual fold change (A) or delta (B) values. Asterisks represent significance of a Wilcoxon rank-sum test: NS. $P \geq 0.05$, * $P < 0.05$, ** $P < 0.01$ , ***$P < 0.001$.

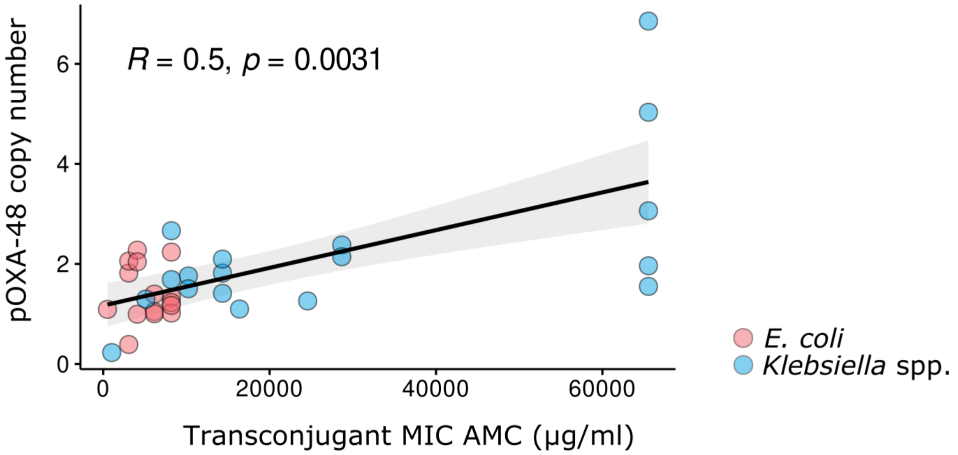


**Fig. S13.** Spearman's correlation between pOXA-48 copy number and the minimum inhibitory concentration (MIC) of amoxicillin/clavulanic acid (AMC) in the pOXA-48-carrying clones. pOXA-48 copy number was estimated from sequencing data as the ratio of plasmid/chromosome median coverage (see Methods).

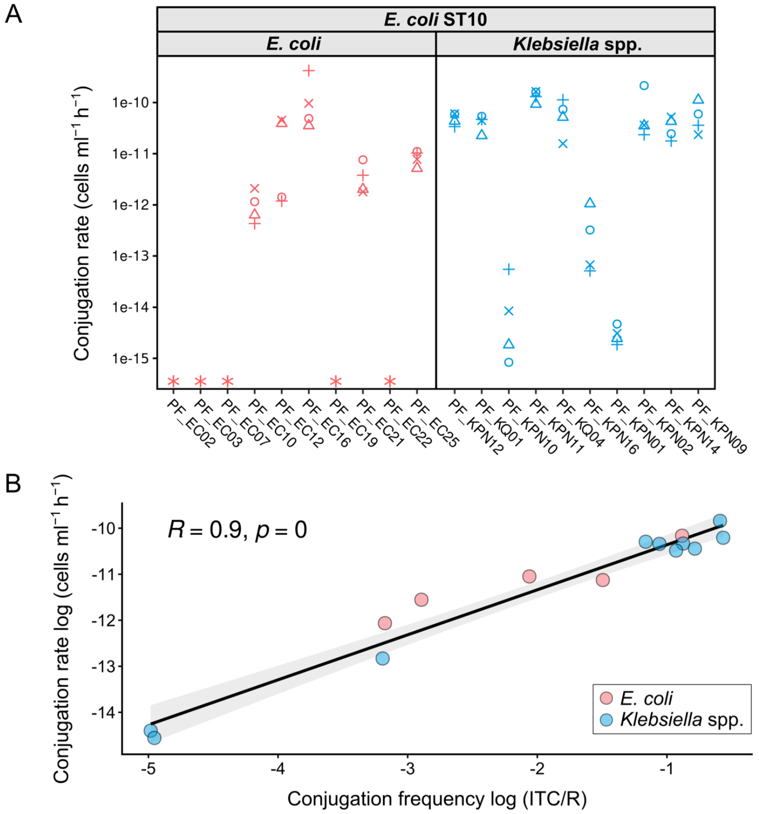


**Fig. S14.** pOXA-48 conjugation rates. We calculated the conjugation rates with *E. coli* ST10 as donor and the 20 different recipient strains. We calculated the rates from the same mating experiments used to calculate pOXA-48 conjugation frequencies. (A) Conjugation rates in cells x ml$^{-1}$ x h$^{-1}$ obtained in conjugation assays with one donor (*E. coli* ST10) and one recipient in equal proportions. Color represents the recipient species (red, *E. coli*; blue, *Klebsiella spp.*), symbols represent biological replicates (n = 4), and asterisks symbolize conjugation frequencies below the detection limit (dotted line). (B) Pearson correlation between pOXA-48 conjugation frequency (TC/R) and rate (cells x ml$^{-1}$ x h$^{-1}$) from mating assays with one donor (*E. coli* ST10) and one recipient in equal proportions. We only used the data from strains with conjugation levels beyond the limit of detection (TC detected in the mating assays).

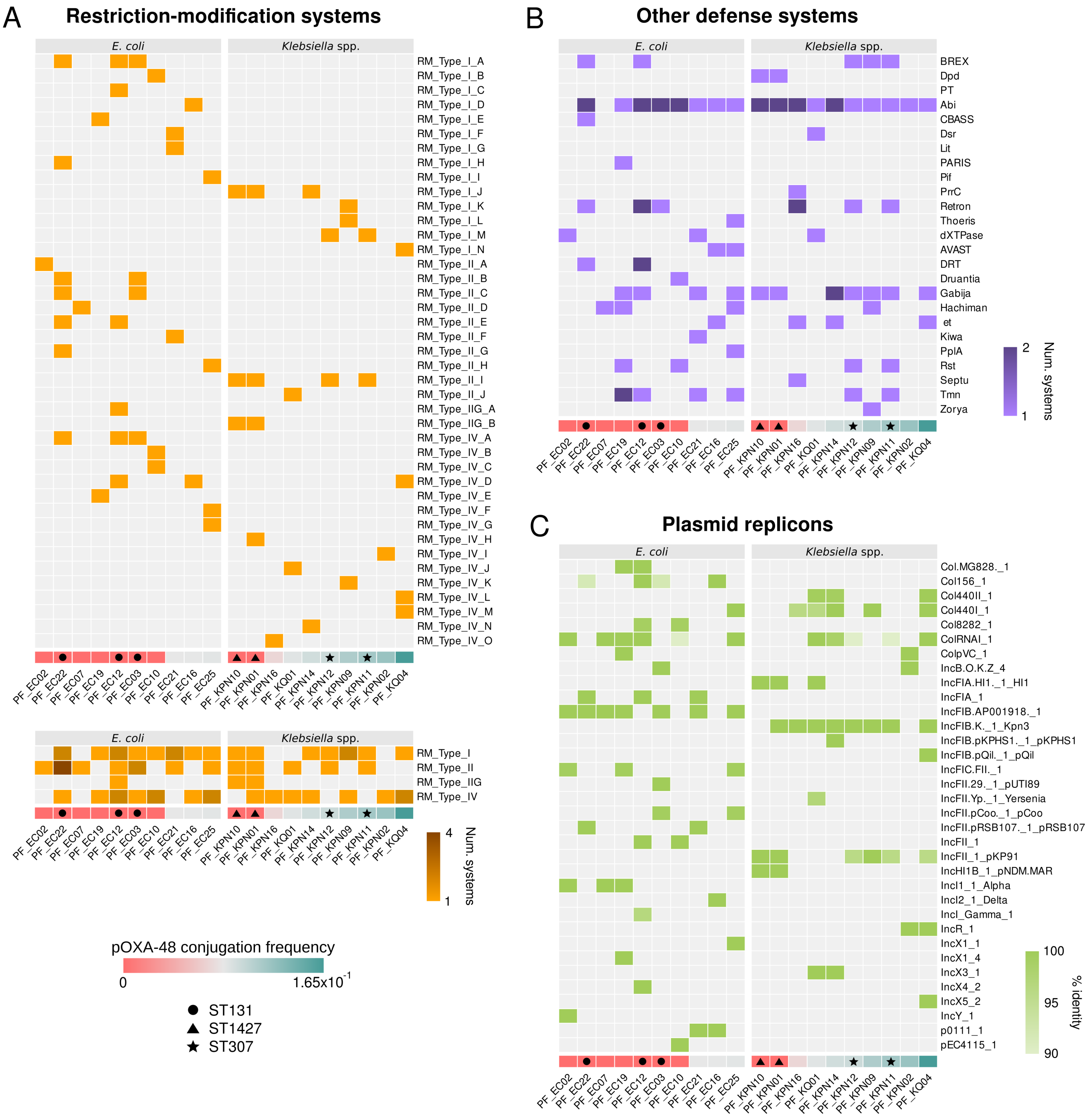


**Fig. S15.** Analysis of genomic traits in recipient strains that could explain differences in pOXA-48 acquisition by conjugation. (A) Heatmap of presence/absence of restriction-modification (RM) systems. In the top panel, all RM systems subtypes, labeled with different letters, are shown. In the bottom panel, RM systems are collapsed by type. (B) Heatmap of presence/absence of other phage defense systems. (C) Heatmap of presence/absence of plasmid replicons. The intensity of the color scale in A bottom and B indicate the number of systems of a specific type a strain is encoding, while in C indicates the percentage of identity of the replicon to the BLAST hit of the PlasmidFinder database (see Methods). In A-C, recipient strains are ordered by mean pOXA-48 conjugation frequencies across replicates and experimental conditions. Strains belonging to the same sequence type (ST) are marked with a symbol.

**Table S1. Model parameters for bacterial strains used in this study.**

| Name | Specie | Type | PCN | MIC | Conjugation rate | Specific affinity | Resource conversion | Resistance coefficient |
|---|---|---|---|---|---|---|---|---|
| PF_EC02 | E | TC | 1.3351 | 8192 | - | $7.60 \times 10^{-10}$ | $5.504 \times 10^{8}$ | $6.25 \times 10^{-2}$ |
| PF_EC03 | E | TC | 2.2371 | 6144 | - | $7.60 \times 10^{-10}$ | $8.90 \times 10^{8}$ | $6.25 \times 10^{-2}$ |
| PF_EC07 | E | TC | 1.0054 | 6144 | - | $3.79 \times 10^{-10}$ | $1.04 \times 10^{9}$ | $6.25 \times 10^{-2}$ |
| PF_EC10 | E | TC | 0.3899 | 3072 | - | $5.66 \times 10^{-10}$ | $1.071 \times 10^{9}$ | $3.125 \times 10^{-2}$ |
| PF_EC12 | E | TC | 1.41 | 6144 | - | $5.19 \times 10^{-10}$ | $9.094 \times 10^{8}$ | $6.25 \times 10^{-2}$ |
| PF_EC16 | E | TC | 1.229 | 8192 | - | $4.91 \times 10^{-10}$ | $1.122 \times 10^{9}$ | $6.25 \times 10^{-2}$ |
| PF_EC19 | E | TC | 1.41 | 6144 | - | $6.08 \times 10^{-10}$ | $9.812 \times 10^{8}$ | $6.25 \times 10^{-2}$ |
| PF_EC21 | E | TC | 1.1712 | 8192 | - | $5.55 \times 10^{-10}$ | $1.021 \times 10^{9}$ | $6.25 \times 10^{-2}$ |
| PF_EC22 | E | TC | 0.9955 | 6144 | - | $6.15 \times 10^{-10}$ | $9.736 \times 10^{8}$ | $6.25 \times 10^{-2}$ |
| PF_EC25 | E | TC | 1.0923 | 512 | - | $6.78 \times 10^{-10}$ | $9.237 \times 10^{8}$ | $3.906 \times 10^{-3}$ |
| PF_KPN01 | K | TC | 1.0987 | 16384 | - | $4.84 \times 10^{-10}$ | $8.746 \times 10^{8}$ | $1.25 \times 10^{-1}$ |
| PF_KPN02 | K | TC | 2.12 | 28672 | - | $6.64 \times 10^{-10}$ | $7.905 \times 10^{8}$ | $2.50 \times 10^{-1}$ |
| PF_KQ01 | K | TC | 2.12 | 65536 | - | $4.67 \times 10^{-10}$ | $1.077 \times 10^{9}$ | $5.00 \times 10^{-1}$ |
| PF_KPN09 | K | TC | 1.7607 | 10240 | - | $6.21 \times 10^{-10}$ | $8.064 \times 10^{8}$ | $1.25 \times 10^{-1}$ |
| PF_KPN10 | K | TC | 2.12 | 65536 | - | $4.83 \times 10^{-10}$ | $7.99 \times 10^{8}$ | $5.00 \times 10^{-1}$ |
| PF_KPN11 | K | TC | 6.8543 | 65536 | - | $5.38 \times 10^{-10}$ | $9.71 \times 10^{8}$ | $5.00 \times 10^{-1}$ |
| PF_KQ04 | K | TC | 1.2603 | 24576 | - | $5.74 \times 10^{-10}$ | $7.568 \times 10^{8}$ | $2.50 \times 10^{-1}$ |
| PF_KPN12 | K | TC | 1.8173 | 14336 | - | $5.44 \times 10^{-10}$ | $9.635 \times 10^{8}$ | $1.25 \times 10^{-1}$ |
| PF_KPN14 | K | TC | 2.12 | 65536 | - | $4.16 \times 10^{-10}$ | $8.994 \times 10^{8}$ | $5.00 \times 10^{-1}$ |
| PF_KPN16 | K | TC | 0.2286 | 1024 | - | $5.39 \times 10^{-10}$ | $8.523 \times 10^{8}$ | $7.812 \times 10^{-3}$ |
| PF_EC02 | E | WT | - | 32 | - | $6.65 \times 10^{-10}$ | $6.786 \times 10^{8}$ | $2.441 \times 10^{-4}$ |
| PF_EC03 | E | WT | - | 16 | - | $6.54 \times 10^{-10}$ | $8.151 \times 10^{8}$ | $1.221 \times 10^{-4}$ |
| PF_EC07 | E | WT | - | 64 | - | $4.15 \times 10^{-10}$ | $1.013 \times 10^{9}$ | $4.883 \times 10^{-4}$ |
| PF_EC10 | E | WT | - | 64 | $8.629 \times 10^{-13}$ | $7.37 \times 10^{-10}$ | $8.472 \times 10^{8}$ | $4.883 \times 10^{-4}$ |
| PF_EC12 | E | WT | - | 64 | $7.48 \times 10^{-12}$ | $4.81 \times 10^{-10}$ | $9.768 \times 10^{8}$ | $4.883 \times 10^{-4}$ |
| PF_EC16 | E | WT | - | 32 | $6.853 \times 10^{-11}$ | $5.55 \times 10^{-10}$ | $1.04 \times 10^{9}$ | $2.441 \times 10^{-4}$ |
| PF_EC19 | E | WT | - | 16 | - | $6.20 \times 10^{-10}$ | $9.424 \times 10^{8}$ | $1.221 \times 10^{-4}$ |
| PF_EC21 | E | WT | - | 64 | $2.79 \times 10^{-12}$ | $4.90 \times 10^{-10}$ | $8.386 \times 10^{8}$ | $4.883 \times 10^{-4}$ |
| PF_EC22 | E | WT | - | 16 | - | $6.62 \times 10^{-10}$ | $9.886 \times 10^{8}$ | $1.221 \times 10^{-4}$ |
| PF_EC25 | E | WT | - | 128 | $8.95 \times 10^{-12}$ | $5.47 \times 10^{-10}$ | $1.004 \times 10^{9}$ | $9.766 \times 10^{-4}$ |
| PF_KPN01 | K | WT | - | 128 | $2.512 \times 10^{-15}$ | $3.65 \times 10^{-10}$ | $8.673 \times 10^{8}$ | $9.766 \times 10^{-4}$ |
| PF_KPN02 | K | WT | - | 64 | $3.595 \times 10^{-11}$ | $6.35 \times 10^{-10}$ | $7.952 \times 10^{8}$ | $4.883 \times 10^{-4}$ |
| PF_KQ01 | K | WT | - | 128 | $4.557 \times 10^{-11}$ | $5.49 \times 10^{-10}$ | $9.948 \times 10^{8}$ | $9.766 \times 10^{-4}$ |
| PF_KPN09 | K | WT | - | 128 | $4.636 \times 10^{-11}$ | $7.30 \times 10^{-10}$ | $7.772 \times 10^{8}$ | $9.766 \times 10^{-4}$ |
| PF_KPN10 | K | WT | - | 64 | $3.981 \times 10^{-15}$ | $4.47 \times 10^{-10}$ | $7.967 \times 10^{8}$ | $4.883 \times 10^{-4}$ |
| PF_KPN11 | K | WT | - | 256 | $1.443 \times 10^{-10}$ | $5.62 \times 10^{-10}$ | $8.49 \times 10^{8}$ | $1.953 \times 10^{-3}$ |
| PF_KQ04 | K | WT | - | 64 | $6.193 \times 10^{-11}$ | $8.67 \times 10^{-10}$ | $6.601 \times 10^{8}$ | $4.883 \times 10^{-4}$ |
| PF_KPN12 | K | WT | - | 128 | $5.073 \times 10^{-11}$ | $6.01 \times 10^{-10}$ | $1.008 \times 10^{9}$ | $9.766 \times 10^{-4}$ |
| PF_KPN14 | K | WT | - | 64 | $3.264 \times 10^{-11}$ | $5.83 \times 10^{-10}$ | $7.306 \times 10^{8}$ | $4.883 \times 10^{-4}$ |
| PF_KPN16 | K | WT | - | 16 | $1.478 \times 10^{-13}$ | $4.36 \times 10^{-10}$ | $8.465 \times 10^{8}$ | $1.221 \times 10^{-4}$ |
